# tRNA modifying enzymes MnmE and MnmG are essential for *Plasmodium falciparum* apicoplast maintenance

**DOI:** 10.1101/2024.12.21.629855

**Authors:** Rubayet Elahi, Luciana Ribeiro Dinis, Russell P. Swift, Hans B. Liu, Sean T. Prigge

**Affiliations:** Department of Molecular Microbiology and Immunology, Johns Hopkins University, Baltimore, Maryland, USA; The Johns Hopkins Malaria Research Institute, Baltimore, Maryland, USA

**Keywords:** Plasmodium, tRNA modification, MnmE, MnmG, apicoplast

## Abstract

The circular genome of the *Plasmodium falciparum* apicoplast contains a complete minimal set of tRNAs, positioning the apicoplast as an ideal model for studying the fundamental factors required for protein translation. Modifications at tRNA wobble base positions, such as xm^5^s^2^U, are critical for accurate protein translation. These modifications are ubiquitously found in tRNAs decoding two-family box codons ending in A or G in prokaryotes and in eukaryotic organelles. Here, we investigated the xm^5^s^2^U biosynthetic pathway in the apicoplast organelle of *P. falciparum*. Through comparative genomics, we identified orthologs of enzymes involved in this process: SufS, MnmA, MnmE, and MnmG. While SufS and MnmA were previously shown to catalyze s^2^U modifications, we now show that MnmE and MnmG are apicoplast-localized and contain features required for xm^5^s^2^U biosynthetic activity. Notably, we found that *P. falciparum* lacks orthologs of MnmC, MnmL, and MnmM, suggesting that the parasites contain a minimal xm^5^s^2^U biosynthetic pathway similar to that found in bacteria with reduced genomes. Deletion of either MnmE or MnmG resulted in apicoplast disruption and parasite death, mimicking the phenotype observed in Δ*mnmA* and Δ*sufS* parasites. Our data strongly support the presence and essentiality of xm^5^s^2^U modifications in apicoplast tRNAs. This study advances our understanding of the minimal requirements for protein translation in the apicoplast organelle.

**Importance:** The apicoplast of *Plasmodium falciparum* is an ideal model for studying minimal translation machinery since it contains the fewest number of tRNA isotypes required for protein translation. In such reduced systems, tRNA modifications, particularly at the wobble anticodon position, are critical for proper decoding of mRNA codons by tRNAs. In this study, we investigated the enzymes responsible for the wobble modification xm^5^s^2^U in the apicoplast. Our findings reveal that only four enzymes are required in the apicoplast to carry out this modification, similar to the situation found in bacteria with highly reduced genomes. Based on their phylogenetic conservation, these four enzymes have been included in all proposed minimal genome concepts to date and it appears that the apicoplast has retained the same core machinery. Our findings provide novel insight into apicoplast biology and into minimal requirements for protein translation.

## Introduction

Transfer RNAs (tRNAs) are small, non-coding RNAs essential for protein translation, converting mRNA codons into amino acids through codon-anticodon interactions on the ribosome (1, 2). The first two codon positions pair stringently with the second and third anticodon bases (positions 36 and 35, respectively) via Watson-Crick base pairing, while the third codon base (wobble position) interacts more flexibly with the first anticodon base (position 34, **Figure 1A**) (2). This flexibility allows a single tRNA to recognize up to four codons.

**Figure 1.**
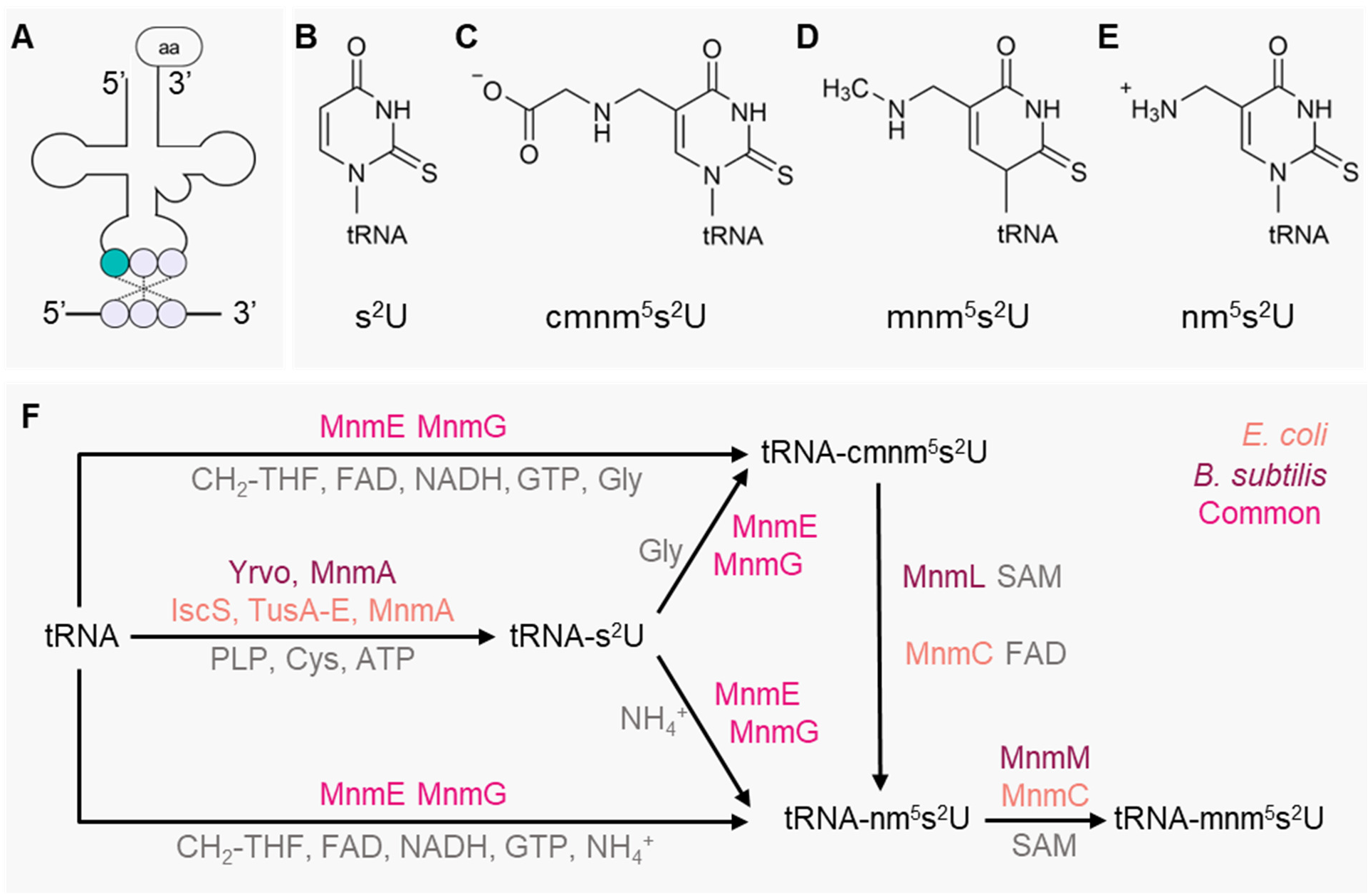
xm^5^s^2^U modification biosynthesis pathway in *Escherichia coli* and *Bacillus subtilis*. (A) Schematic representation of codon-anticodon recognition during protein translation. The tRNA anticodon loop (highlighted with the wobble position in teal) pairs with the mRNA codon, while the amino acid (aa) is attached at the CCA terminus. (B-E) Chemical structures of tRNA wobble position modifications: (B) s^2^U modification at the C2 position, (C) carboxymethyl-amino (cmnm^5^), (D) methyl-amino (mnm^5^), and (E) amino (nm^5^) modifications at the C5 position. (F) Comparison between the xm^5^s^2^U modification pathways in model bacterial systems. Enzymes specific to *Escherichia coli* (orange), *Bacillus subtilis* (purple), and those common to both organisms (magenta) are shown. The MnmE-MnmG complex functions either independently to install xm^5^U modifications in tRNAs or modifies pre-existing s^2^U bases to generate xm^5^s^2^U. The s^2^U modifications are installed by organism-specific pathways: YrvO and MnmA in *B. subtilis*, or IscS, TusA-E and MnmA in *E. coli*. Final modifications are also organism-specific: *E. coli* uses MnmC, while *B. subtilis* employs MnmL and MnmM for terminal modifications. Substrates and cofactors are shown in gray.

Wobble base flexibility is potentially problematic for coding amino acids that share the same codon box (such as histidine and glutamine codons which differ only at the wobble base position). Specificity in these cases is achieved through tRNA modifications across all kingdoms (3), with over 30 distinct types of wobble modification identified to date (4). These modifications fine-tune wobble base pairing, allowing a single tRNA to accurately decode codons differing only in the third nucleotide. For instance, tRNAs decoding two-family box codons ending in A or G often carry xm^5^s^2^U modifications (5). These modifications restrict the wobble capacity of uridine (U34) by introducing a thiolation at position 2 (C2) of U34 (s^2^U) (**Figure 1B**), which stabilizes the C3′-endo conformation of the ribose. This structural shift improves base pairing with purines (A or G) and prevents misreading U-or C-ending codons (6–8). The xm^5^U modification at position 5 (C5) of U34 enhances U•G pairing and improves reading accuracy for NNA codons (9, 10). In these modifications, “x” represents carboxymethyl-amino (cmn, **Figure 1C**), methyl-amino (mn, **Figure 1D**), or amino (n, **Figure 1E**) moieties.

The biosynthesis of xm^5^s^2^U modifications is intricate, with xm^5^ and s^2^ moieties added independently through two distinct pathways, each involving multiple enzymes (5, 9, 11–13). It has been suggested that the s^2^U modification can precede xm^5^U modification (13, 14). In gram-negative model bacterium *Escherichia coli*, the s^2^U biosynthesis pathway involves seven enzymes: IscS, TusA-E, and MnmA. The pyridoxal phosphate (PLP)-dependent IscS liberates sulfur from cysteine, which is subsequently transferred to MnmA via the sulfurtransferases TusA-E. MnmA then incorporates this sulfur into the C2 position of U34 in tRNAs (tRNA^Lys^, tRNA^Glu^, and tRNA^Gln^) (**Figure 1F**) (6, 7, 15). Conversely, in the gram-positive bacterium *Bacillus subtilis* s^2^U biosynthesis relies on the PLP-dependent cysteine desulfurase YrvO, which directly transfers sulfur from cysteine to MnmA (**Figure 1F**) (6).

In both *E. coli* and *B. subtills*, the xm^5^U modifications are installed by the MnmE-MnmG enzyme complex, which can generate either cmnm^5^U or nm^5^U derivatives depending on whether glycine or ammonium (NH_4_^+^) serves as the substrate (16). MnmE functions as a homodimer, with each monomer containing three domains: an N-terminal dimerization domain that binds methylene-tetrahydrofolate (CH_2_-THF), a central helical domain, and a discrete G domain that binds guanosine 5’-triphosphate (GTP) (17–20).

Its partner protein, MnmG, also forms a homodimer and comprises a flavin adenine dinucleotide (FAD)-binding domain, two insertion domains, and a C-terminal helical domain that mediates MnmE interaction (21–24). The complete modification reaction requires three cofactors: GTP, FAD, NADH, and CH_2_-THF, with the latter providing the methylene group for C5 modification (11, 17, 21, 25–28) (Figure 1F). In bacteria like E. coli, the cmnm^5^U derivative can be further modified into mnm^5^U by the bifunctional enzyme MnmC (29), which contains two distinct catalytic domains. The C-terminal domain (MnmC1) facilitates the FAD-dependent oxidoreduction of cmnm^5^s^2^U to nm^5^s^2^U. The N-terminal domain (MnmC2) catalyzes the methylation of nm^5^s^2^U to mnm^5^s^2^U, using S-Adenosyl-*L*-Methionine (SAM) as the methyl group donor (29). However, in *B. subtills*, MnmL facilitates conversion of cmnm^5^s^2^U to nm^5^s^2^U in a SAM dependent manner (14) and MnmM catalyzes the methylation of nm^5^s^2^U to mnm^5^s^2^U using SAM as the methyl group donor (30) (Figure 1F). These modifications occur at the C5 of U34 in tRNA_Lys_, tRNA_Glu_, tRNA_Gln_, tRNA_Leu_, and tRNA_Arg_ (5, 9, 17, 23, 25, 31).

The apicoplast, a relict plastid in the apicomplexan parasite *Plasmodium falciparum* encodes only 25 tRNA isotypes in its genome, the smallest known number for any translational machinery (32–39). This reduced tRNA diversity, combined with the lack of evidence for tRNA import into secondary plastids like *P. falciparum* apicoplast (39), makes the apicoplast an ideal system for investigating the minimal requirements of translational machinery. tRNA modifications, like xm^5^s^2^U, are vital for maintaining the accuracy and efficiency of protein translation. Given the prokaryotic origin of apicoplast and the ubiquitous occurrence of xm^5^s^2^U modifications in prokaryotes, we hypothesize that such modifications exist in apicoplast tRNAs. Previous studies have demonstrated the importance of SufS-MnmA-mediated s^2^U modifications for protein translation in the apicoplast (40). However, the xm^5^s^2^U biosynthetic pathway in the *P. falciparum* apicoplast remains unexplored.

The apicoplast organellar genome contains a full set of tRNAs but does not encode any putative tRNA modification enzymes. These enzymes must be among the hundreds of nucleus-encoded proteins that are trafficked to the apicoplast using N-terminal leader sequences (41). In this study, we applied comparative genomics to identify the orthologs of the enzymes involved in xm^5^s^2^U biosynthesis and showed that these nucleus-encoded proteins are trafficked to the apicoplast. Using an apicoplast metabolic bypass system (42), we demonstrated that these enzymes are crucial for apicoplast maintenance and parasite survival. These findings offer additional insights into the minimal requirements for protein translation, marking the first characterization of the xm^5^s^2^U biosynthesis pathway in an apicomplexan parasite. This study significantly advances our understanding of apicoplast biology and the broader biology of the parasite.

## Results

### The apicoplast of *Plasmodium falciparum* contains a reduced xm^5^s^2^U tRNA modification pathway

The prokaryotic origin of the *P. falciparum* apicoplast (33, 34, 36, 38) prompted us to hypothesize that it may harbor an xm^5^s^2^U biosynthesis pathway analogous to those found in bacterial systems. To investigate this hypothesis, we employed enzymes from both gram-negative model bacteria *E. coli* and gram-positive model bacteria *B. subtilis*, known to be involved in the xm^5^s^2^U biosynthesis pathway, as queries to search for orthologs within the *P. falciparum* genome (**Supplementary Table 1**). We identified orthologs for five proteins in *Plasmodium*, each sharing 24 to 50% sequence identity with their bacterial counterparts (**Table 1**, **Supplementary Table 2**). Notably, these five orthologs were also found in other pathogenic apicomplexans (**Table 1, Supplementary Table 2**). Importantly, all identified *P. falciparum* orthologs are encoded by the nuclear genome. The apicoplast imports these proteins via the secretory pathway, mediated by an N-terminal leader peptide which is necessary and sufficient for apicoplast trafficking (41). Multiple bioinformatic tools are available to predict the presence of these leader peptides and to evaluate the likelihood of apicoplast localization. We used PlasmoAP (43), PATS (44), and ApicoAP (45) to predict the localization of the *Plasmodium* orthologs involved in the xm^5^s^2^U biosynthesis pathway. Additionally, we cross-referenced our predictions with data from the recently published apicoplast proteome, which was identified using proximity biotinylation-based proteomics (BioID) (46). Of the five *P. falciparum* proteins identified, four were predicted to localize to the apicoplast by at least two bioinformatic methods (**Table 2**). Although IscS was predicted by one tool to contain an apicoplast targeting signal, neither the other prediction tools nor the BioID-derived apicoplast proteome identified this protein as part of the apicoplast proteome. Furthermore, IscS has been previously shown to localize to the mitochondrion (47).

**Table 1.** List of xm^5^s^2^U biosynthesis pathway enzyme orthologs of pathogenic apicomplexans identified by homology search with bacterial enzymes as queries. Orthologous enzyme gene identifiers are from ^a^PlasmoDB, ^b^ToxoDB, ^c^PiroplasmaDB, and ^d^CryptoDB.-, not detected.

| Enzyme | *P. falciparum*<br>(PF3D7_)<sup>a</sup> | *T. gondii*<br>(TGME49_)<sup>b</sup> | *E. tenella*<br>(ETH2_)<sup>b</sup> | *B. microti*<br>(BMR1_)<sup>c</sup> | *T. annulata*<br>(TA)<sup>c</sup> | *C. parvum*<br>(cgd4_)<sup>d</sup> |
| --- | --- | --- | --- | --- | --- | --- |
| IscS | 0727200 | 211090 | 1354600 | 03g04230 | 19715 | 3040 |
| YrvO | 0716600 | 216170 | 1112500 | 03g01415 | 16295 | - |
| TusA | - | - | - | - | - | - |
| TusB | - | - | - | - | - | - |
| TusC | - | - | - | - | - | - |
| TusD | - | - | - | - | - | - |
| TusE | - | - | - | - | - | - |
| MnmA | 019800 | 309110 | 349800 | 03g03155 | 12620 | - |
| MnmC | - | - | - | - | - | - |
| MnmE | 0817100 | 204350 | 1106900 | 02g02630 | 17200 | - |
| MnmG | 1244000 | 242010 | 1518800 | 04g09035 | 09230 | - |
| MnmL | - | - | - | - | - | - |
| MnmM | - | - | - | - | - | - |

**Table 2.** Predicted and experimental localization of *Plasmodium falciparum* xm^5^s^2^U modification biosynthesis enzymes. The likelihood of apicoplast localization for each enzyme is indicated as either Yes (Y) or No (N). ^a^PlasmoDB, ^b^Foth *et al*., 2003 [43]; ^c^Cilingir *et al*., 2012 [45]; ^d^Zuegge *et al*., 2001 [44]; ^e^Boucher *et a*l., 2018 [46]; ^1^Gisselberg *et al.*, 2013 [47]; ^2^Swift, Elahi *et al*., 2023 [40].

| Gene ID<br>(PF3D7_)<sup>a</sup> | Name | PlasmoAP<sup>b</sup> | ApicoAP<sup>c</sup> | PATS<sup>d</sup> | BiolD<sup>e</sup> | Experimental<br>localization |
| --- | --- | --- | --- | --- | --- | --- |
| 0727200 | IscS | Y | N | N | N | Mitochondrion<sup>1</sup> |
| 0716600 | SufS | Y | Y | Y | Y | Apicoplast<sup>1</sup> |
| 1019800 | MnmA | N | Y | Y | N | Apicoplast<sup>2</sup> |
| 0817100 | MnmE | Y | Y | Y | N | ? |
| 1244000 | MnmG | N | Y | Y | Y | ? |

Among the remaining four proteins, SufS and MnmA have been previously confirmed by our group to localize to the apicoplast (40), consistent with predictions by at least two bioinformatic tools (**Table 2**). SufS is a cysteine desulfurase that was initially thought to function exclusively within the apicoplast-localized SUF iron-sulfur (FeS) cluster pathway (47, 48). We recently showed that SufS has an additional role in supplying sulfur to MnmA for the s^2^U modification of apicoplast tRNAs (40).

Complementation of parasite MnmA and/or SufS with *B. subtilis* MnmA and *B. subtilis* cysteine desulfurase YrvO supported a model of direct sulfur transfer from the cysteine desulfurase to MnmA without a requirement for other sulfur transfer proteins (40).

The final two proteins we identified appeared to be *P. falciparum* orthologs of bacterial MnmE and MnmG enzymes. MnmE and MnmG are homodimeric proteins that form an α_2_β_2_ and α_4_β_2_ complexes which catalyzes the cmnm^5^ or nm^5^ modification at C5 of U34 in substrate tRNAs (11, 17, 21, 25–28, 49). The *Plasmodium* ortholog of MnmE is annotated as a ‘tRNA modification GTPase’ and the MnmG ortholog is annotated as a ‘glucose-inhibited division protein A homologue’ (35). Hereafter, we referred to them as MnmE and MnmG, due to the strong conservation of functionally important residues and structural homology with bacterial counterparts, as discussed below.

Multiple sequence alignment of the putative *P. falciparum* MnmE with orthologs from various organisms, including *E. coli*, *B. subtilis*, *Pseudomonas syringae*, *Thermotoga maritima*, *Saccharomyces cerevisiae*, *Arabidopsis thaliana*, and *Homo sapiens*, revealed an N-terminal extension of about 160 amino acids (Figure 2A, **Supplementary Figure 1**). This extension is predicted to serve as a leader peptide, directing the protein to the apicoplast via the secretory pathway. The 98 kDa *P. falciparum* MnmE retains all conserved residues required for CH_2_-THF binding and the GTP-binding motifs (G1-G4) essential for its functionality (Figure 2A, **Supplementary Figure 1**). Additionally, the highly conserved CxGK tetrapeptide motif at the C-terminus of the enzyme, which is crucial for tRNA binding and modification activity among MnmE orthologs (17), is also present in *P. falciparum* MnmE (**Supplementary Figure 1**). To gain deeper structural insights into *P. falciparum* MnmE, we used AlphaFold (50–52) to predict its three-dimensional structure. Our analysis revealed a predicted folding architecture that resembles the structure of *Thermotoga maritima* MnmE (*Tm*MnmE) (20) (Figure 2B). The structural analysis of *P. falciparum* MnmE shows the presence of all the canonical domains found in MnmE orthologs from other organisms: a N-terminal α/β domain responsible for homodimerization and CH_2_-THF binding, a central helical domain, and a G domain with a Ras-like fold crucial for GTP binding. Moreover, a structural alignment of the predicted *P. falciparum* MnmE with the crystal structure of *Tm*MnmE demonstrated a high degree of similarity, with a root mean square deviation (RMSD) value of 1.3 Å, indicating a strong agreement between the predicted and experimentally determined structures. Taken together, structural and sequence features of *P. falciparum* MnmE are consistent with those of other MnmE enzymes.

**Figure 2.**
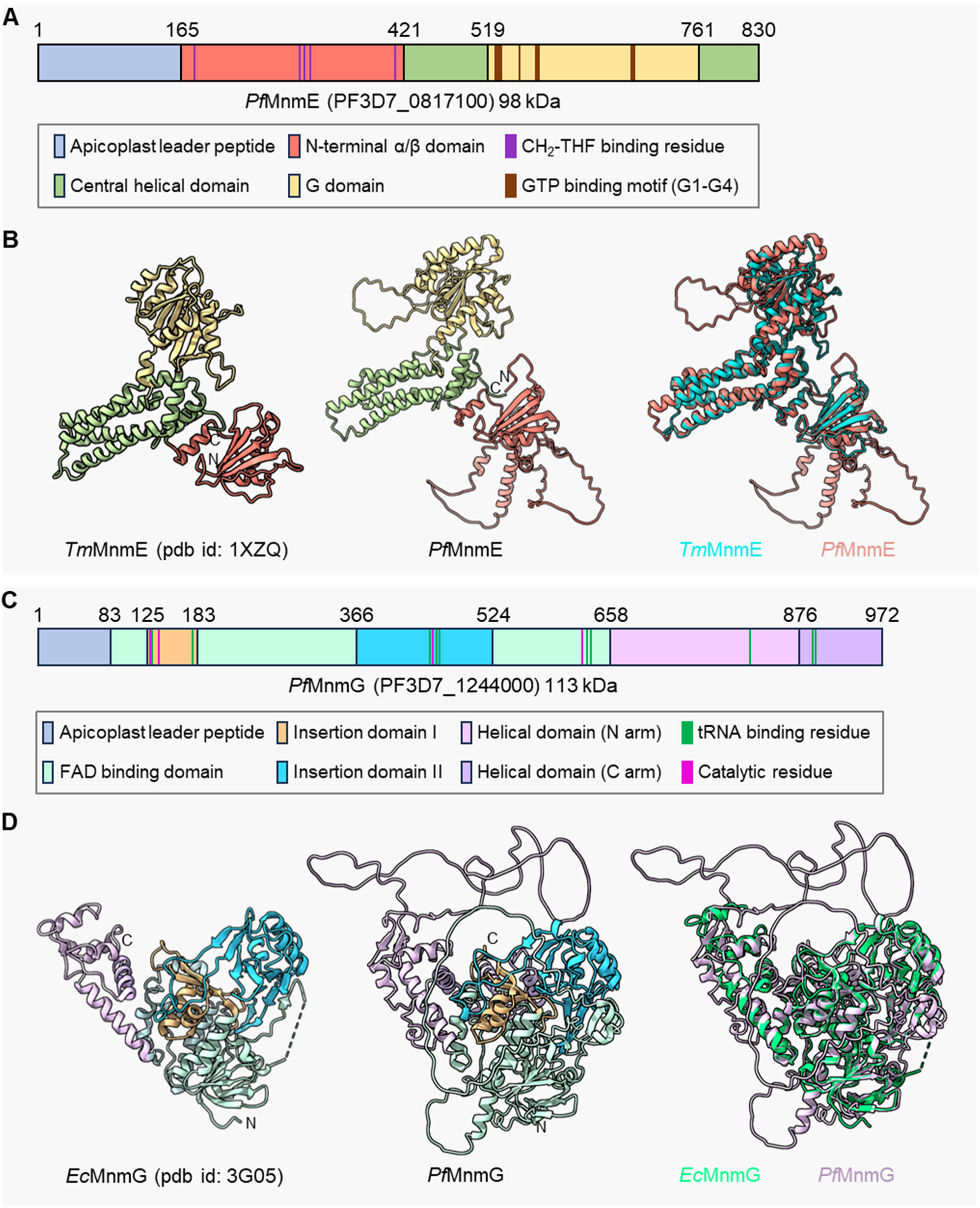
*Plasmodium falciparum* contains MnmE and MnmG orthologs. (A) Domain organization of *P. falciparum* MnmE (*Pf*MnmE, PF3D7_0817100), showing the apicoplast leader peptide, N-terminal α/β domain, CH_2_-THF binding residue, central helical domain, G domain, and GTP binding motifs (G1-G4). (B) Left panel, crystal structure of *Thermotoga maritima* MnmE (*Tm*MnmE, PDB: 1XZQ); middle panel, predicted structure of P*. falciparum* MnmE residues 166-830 (*Pf*MnmE); right panel, structural alignment of *Tm*MnmE (cyan) and *Pf*MnmE (salmon) showing conservation of domain architecture. (C) Domain organization of *P. falciparum* MnmG (*Pf*MnmG, PF3D7_1244000), depicting the apicoplast leader peptide, FAD binding domain, insertion domains I and II, helical domains (N and C arms), tRNA binding residues, and catalytic residues. (D) Left panel, crystal structure of *E. coli* MnmG (*Ec*MnmG, PDB: 3G05); middle panel, predicted structure *of P. falciparum* MnmG *(Pf*MnmG); right panel, structural alignment of *Ec*MnmG (green) and *Pf*MnmG (purple) demonstrating structural conservation. Domain color coding is consistent within panels (A-B) and (C-D).

Similarly, alignment of the putative *P. falciparum* MnmG with orthologs from *E. coli*, *B. subtilis*, *T. maritima*, *Chlorobaculum tepidum*, *A. thaliana*, *S. cerevisiae*, and *H. sapiens* revealed an N-terminal extension of about 80 amino acids (Figure 2C, **Supplementary Figure 2**), likely containing a leader peptide that directs the protein to the apicoplast via the secretory pathway. The 113 kDa *P. falciparum* MnmG retains all conserved catalytic residues predicted for its enzymatic activity (22, 24). Moreover, while most of the conserved residues important for tRNA binding are maintained in *P. falciparum* MnmG, one of these residues is replaced by a similarly charged amino acid (Figure 2C, **Supplementary Figure 2**). This substitution likely preserves a comparable electrostatic potential at the tRNA binding site, potentially allowing for analogous tRNA interactions despite the residue variation. The AlphaFold-predicted structure for *P. falciparum* MnmG exhibited a characteristic funnel-shaped folding architecture, consistent with the experimentally determined structure of *Ec*MnmG (22) (Figure 2D), albeit with low-complexity loop regions in *P. falciparum* MnmG. Structural analysis also revealed a conserved domain organization, including an FAD-binding domain interspersed with two insertion domains (Insertion domain I and II), and a C-terminal helical domain further subdivided into N-arm and C-arm regions, based on sequence and structural similarities (25). A structural alignment of *P. falciparum* MnmG with *Ec*MnmG demonstrated a high degree of similarity, with an RMSD value of 0.9 Å, suggesting that *P. falciparum* MnmG retains the essential functional domains and residues crucial for tRNA modification. Altogether, the *P. falciparum* apicoplast contains only MnmA, MnmE, and MnmG, along with the cysteine desulfurase SufS for xm^5^s^2^U biosynthesis.

### MnmE and MnmG localize to the apicoplast

We previously determined the apicoplast localization of SufS and MnmA and their involvement in s^2^U tRNA modification (40). To validate the predicted localization of MnmE and MnmG, we initially attempted to tag the endogenous proteins at their C-termini with epitope tags. However, multiple efforts to generate these parasite lines were unsuccessful. It is likely that, similar to findings in *E. coli* (17, 18, 21, 25), the C-terminal regions of the *P. falciparum* orthologs are critical for complex formation and enzymatic activity. As an alternative strategy, we fused the predicted N-terminal leader peptides— believed to contain bipartite apicoplast trafficking determinants—of *P. falciparum* MnmE and MnmG to mCherry fluorescent protein. These constructs were expressed from the nuclear genome, allowing us to track their subcellular localization (**Supplementary Figure 3**).

For MnmE, we generated a construct by fusing the leader peptide, consisting of the first 165 N-terminal amino acid residues, to the mCherry protein (designated as *mnmE_lp_*^+^, Figure 3A). This construct was integrated into the genome of PfMev^attB^ parasites (53) using a knock-in approach mediated by mycobacteriophage integrase-based recombination (54) (**Supplementary Figure 3**). The PfMev^attB^ parasite line expresses super-folder green fluorescent protein in the apicoplast (api-SFG) (55), which enables visualization of the apicoplast via live epifluorescence microscopy (42, 53).

**Figure 3.**
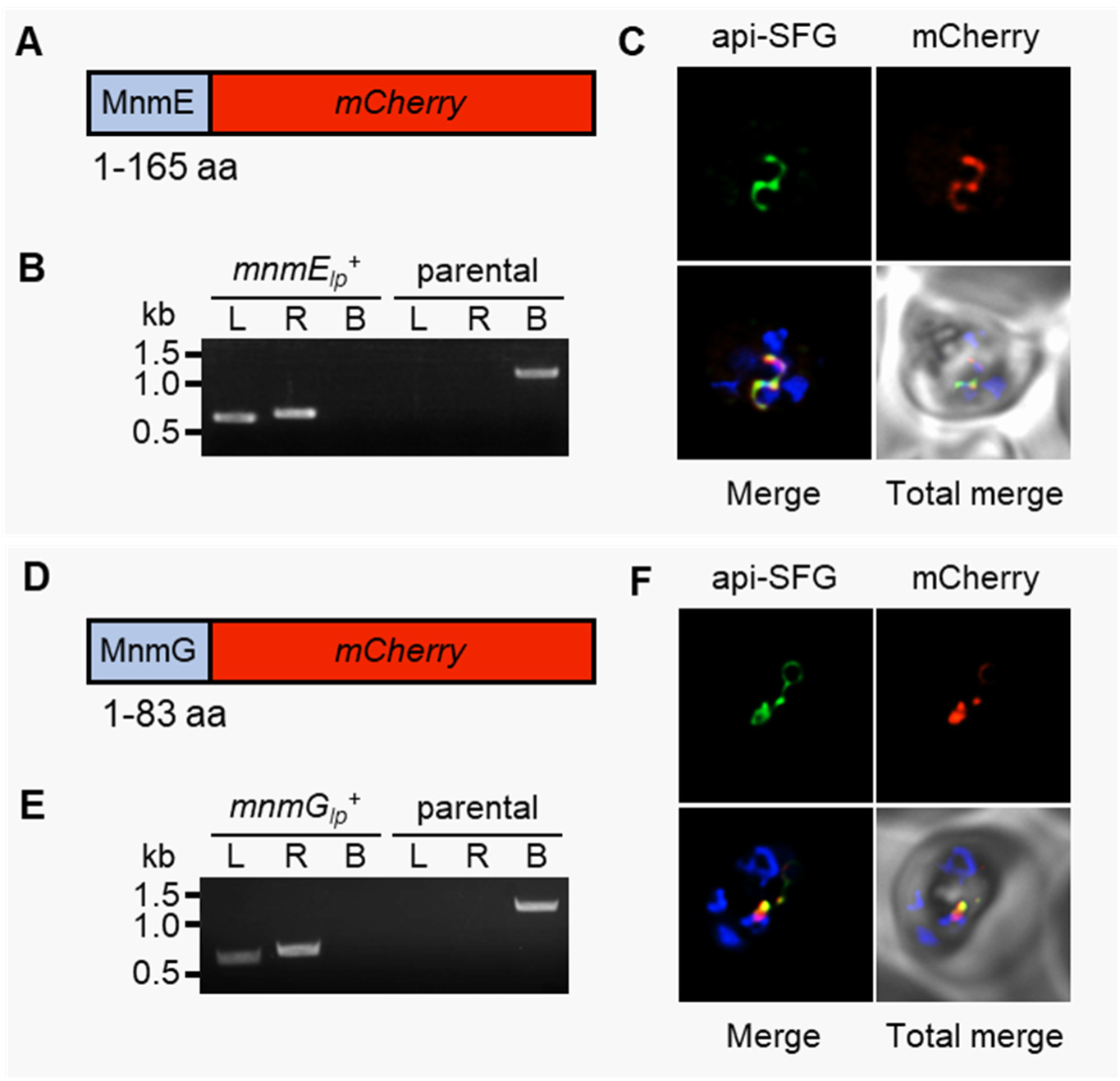
*Plasmodium falciparum* MnmE and MnmG localize to the apicoplast. (A) Schematic representation of the synthetic construct containing the predicted MnmE leader peptide (amino acids 1-165) fused to mCherry fluorescent protein. (B) PCR confirmation of plasmid integration in *mnmE_lp_*^+^ parasites via amplification of attL (L) and attR (R) regions. attB (B) amplification from the PfMev^attB^ parental line serves as control. (C) Representative epifluorescence microscopy images of *mnmE_lp_*^+^ parasites showing colocalization of MnmE_lp_-mCherry fusion protein (red) with apicoplast api-SFG marker (green). (D) Schematic representation of synthetic construct containing the predicted MnmG leader peptide (amino acids 1-83) fused to mCherry fluorescent protein. (E) PCR confirmation of plasmid integration in *mnmG_lp_*^+^ parasites via amplification of attL (L) and attR (R) regions. attB (B) amplification from the PfMev^attB^ parental line serves as control. (F) Representative epifluorescence microscopy images of *mnmG_lp_*^+^ parasites showing colocalization of MnmG_lp_-mCherry fusion protein (red) with apicoplast api-SFG marker (green). DNA size markers in panels (B) and (E) are shown in kilobases (kb). Expected amplicon sizes for panels (B) and (E) and corresponding primer pairs are depicted in **Supplementary** Figure 3. Nuclear DNA in panels (C) and (F) is stained with DAPI (blue). The images in panels (C) and (F) are each 10 μm × 10 μm.

Successful generation of *mnmE*_*lp*_^+^ parasites was confirmed by genotyping PCR (Figure 3B). Live epifluorescence microscopy of *mnmE*_*lp*_^+^ parasites demonstrated colocalization of mCherry fluorescence with apicoplast SFG fluorescence (Figure 3C), indicating that MnmE localizes to the apicoplast.

Similarly, for MnmG, we fused the first 83 N-terminal residues to mCherry (designated as *mnmG_lp_*^+^, Figure 3D) and integrated this construct into PfMev^attB^ parasites (**Supplementary Figure 3**). Genotyping PCR confirmed successful integration (Figure 3E), and live microscopy demonstrated colocalization of mCherry with api-SFG fluorescence (Figure 3F), demonstrating apicoplast localization of MnmG. Collectively, these results show that the amino-terminal regions of both MnmE and MnmG function as apicoplast trafficking peptides capable of directing both proteins to the organelle.

### MnmE and MnmG are essential for apicoplast maintenance

We investigated the essentiality of MnmE and MnmG using CRISPR/Cas9-mediated gene deletion (**Supplementary Figure 4**) in the PfMev parasite line (42). In addition to expressing api-SFG, this parasite line has an engineered cytosolic pathway that synthesizes the isoprenoid precursors IPP (isopentenyl pyrophosphate) and DMAPP (dimethylallyl pyrophosphate) from mevalonate, enabling the parasites to survive without the apicoplast organelle and its endogenous isoprenoid biosynthesis pathway (42) (Figure 4A).

**Figure 4.**
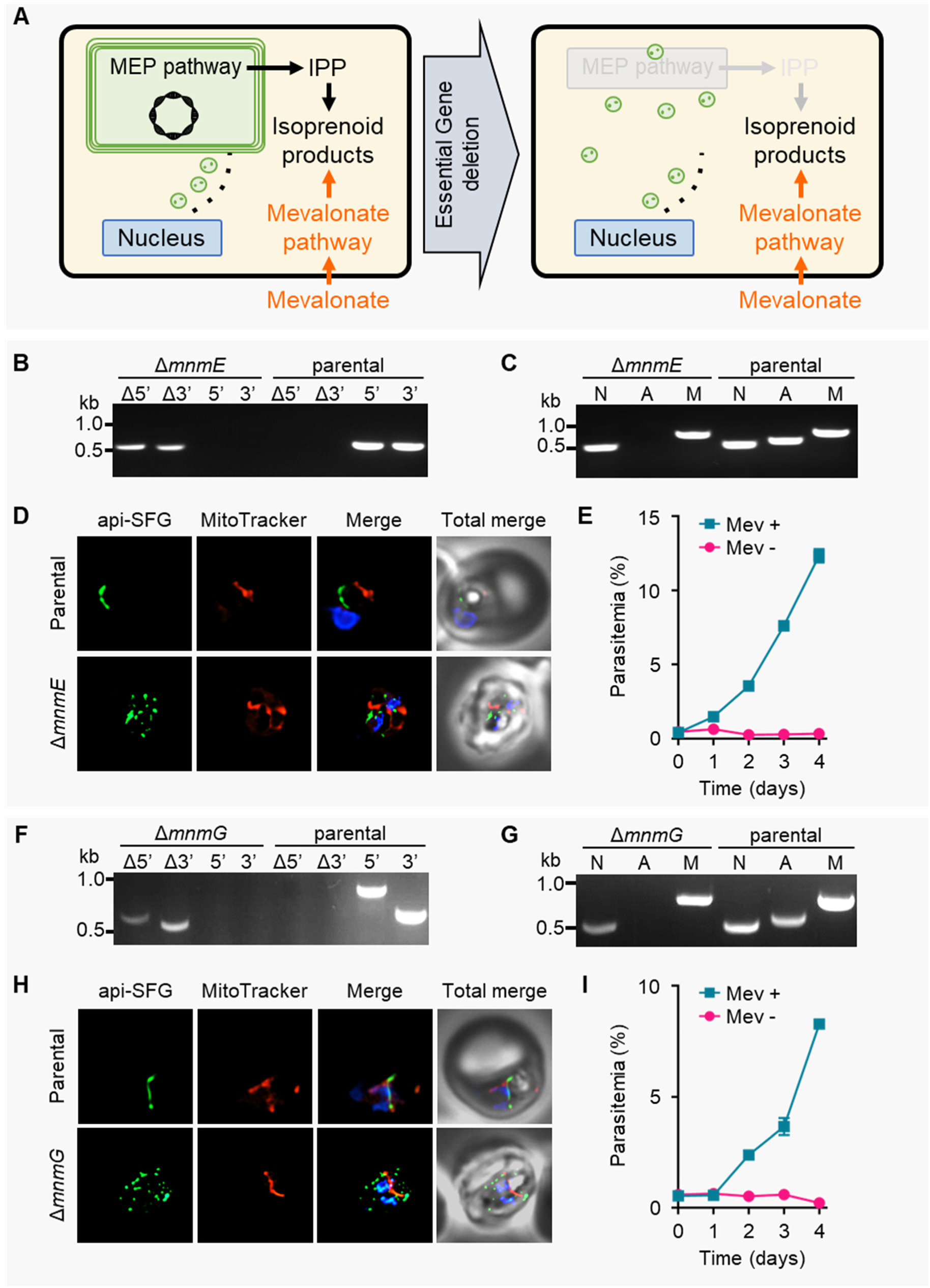
*Plasmodium falciparum* MnmE and MnmG are essential for apicoplast maintenance and parasite survival. (A) PfMev parasites provided with exogenous mevalonate survive loss of genes essential for apicoplast maintenance. Disrupted apicoplasts lose the organellar genome and multiple discrete vesicles containing nucleus-encoded proteins remain. (B) Genotyping PCR verifies *mnmE* deletion in Δ*mnmE* parasites, shown by the presence of amplicons for the Δ5’ and Δ3’ loci at the integration site, but not for the native loci (5’ and 3’) found in the PfMev (parental) parasites. The primers and expected amplicon sizes are depicted in **Supplementary** Figure 4. (C) Attempted PCR amplification of *ldh*, *sufB*, and *cox1* genes of the parasite nuclear (N), apicoplast (A), and mitochondrial (M) genomes, respectively, in Δ*mnmE* and PfMev (parental) parasites. Lack of an amplicon for *sufB* in the Δ*mnmE* parasites indicates the loss of the apicoplast genome. Refer to **Supplementary Table 3** for primer names and sequences and Materials and Methods for expected amplicon sizes. (D) Representative epifluorescence microscopy images of Δ*mnmE* parasites shows multiple discrete vesicles (bottom panel) demonstrating a disrupted apicoplast, compared to an intact apicoplast in PfMev parasites (parental, top panel). Api-SFG protein (green) marks the apicoplast, the mitochondrion is stained with MitoTracker (red), and nuclear DNA is stained with DAPI (blue). (E) The Δ*mnmE* parasites are dependent on mevalonate (Mev) for growth. Asynchronous parasites were grown with or without 50 μM Mev and parasitemia was monitored every 24 h by flow cytometry for 4 days. Data points represent daily mean parasitemia ± standard error of mean (SEM) from two independent biological replicates, each with four technical replicates. (F) Genotyping PCR verifies *mnmG* deletion in Δ*mnmG* parasites, shown by the presence of amplicons for the Δ5’ and Δ3’ loci at the integration site, but not for the native loci (5’ and 3’) found in the PfMev (parental) parasites. The primers and expected amplicon sizes are depicted in Su**pplementary** Figure 4. (G) Attempted PCR amplification of *ldh*, *sufB*, and *cox1* genes of the parasite nuclear (N), apicoplast (A), and mitochondrial (M) genomes, respectively, in Δ*mnmG* and PfMev (parental) parasites. Lack of an amplicon for *sufB* in the Δ*mnmG* parasites indicates the loss of the apicoplast genome. Refer to **Supplementary Table 3** for primer names and sequences and Materials and Methods for expected amplicon sizes. (H) Representative epifluorescence microscopy images of Δ*mnmG* parasites shows multiple discrete vesicles (bottom panel) demonstrating a disrupted apicoplast, compared to an intact apicoplast in PfMev parasites (parental, top panel). Api-SFG protein (green) marks the apicoplast, the mitochondrion is stained with MitoTracker (red), and nuclear DNA is stained with DAPI (blue). (I) The Δ*mnmG* parasites are dependent on mevalonate (Mev) for growth. Asynchronous parasites were grown with or without 50 μM Mev and parasitemia was monitored every 24 h by flow cytometry for 4 days. Data points represent daily mean parasitemia ± standard error of mean (SEM) from two independent biological replicates, each with four technical replicates. DNA markers in panels (B), (C), (F), and (G) are in kilobases (kb). Each image depicts a field of 10 μm ×10 μm in panels (D) and (H).

We confirmed the successful deletion of *mnmE* through genotyping PCR (Figure 4B). Previously, we demonstrated that deletion of s^2^U modification catalyzing enzyme *mnmA* resulted in apicoplast disruption and loss of the organellar genome (40). To determine if *mnmE* deletion similarly affects the apicoplast, we attempted to amplify the apicoplast-encoded gene *sufB* from Δ*mnmE* parasites. These amplification attempts were unsuccessful (Figure 4C), indicating a loss of the apicoplast genome. Consistent with this result, Δ*mnmE* parasites exhibited the classic phenotype of apicoplast disruption—multiple discrete api-SFG-labeled vesicles (56) (Figure 4D). As expected for parasites with a disrupted apicoplast, Δ*mnmE* parasites were strictly dependent on mevalonate for survival (Figure 4E). Similarly, deletion of *mnmG* (Figure 4F) led to apicoplast genome loss (Figure 4G), apicoplast disruption (Figure 4H), and a dependence on mevalonate for the survival of Δ*mnmG* parasites (Figure 4I).

Together, these findings demonstrate that MnmE and MnmG are critical for both parasite survival and apicoplast maintenance. Since both enzymes are predicted to form a heteromeric complex to catalyze essential tRNA modifications, their deletion likely impairs the translation of apicoplast-encoded proteins, leading to apicoplast dysfunction and parasite death.

## Discussion

Modifications in the tRNA anticodon loop, particularly at position 34 (the wobble position), are essential for ensuring translational efficiency and fidelity. This wobble nucleotide interacts with the third base of mRNA codons, with its pairing often modulated by intricate post-transcriptional modifications. The xm^5^s^2^U modification (where x = cmnm^5^, nm^5^, or mnm^5^) exemplifies this complexity in bacterial, chloroplast, and mitochondrial tRNAs by incorporating a sulfur atom (s^2^) and a methyl group with variable side chains (xm^5^) (9, 11, 12, 57, 58). These modifications allow for accurate codon-anticodon interaction and reduction in +1 and +2 frameshifts (6, 8, 31). The xm^5^U modification occurs at C5 of U34 in tRNA_Lys_, tRNA_Glu_, tRNA_Gln_, tRNA_Leu_, and tRNA_Arg_.

For each of these tRNAs the corresponding codons are part of a ‘two-family’ codon box in which two amino acids are encoded and codons ending in A or G must be distinguished from those ending in U or C. In tRNA_Lys_, tRNA_Glu_, and tRNA_Gln_, the xm^5^ addition can be typically preceded by an s^2^U modification at C2 of U34 (13, 14) leading to xm^5^s^2^U modifications. Together, these modifications fine-tune codon recognition by modulating base-pairing flexibility at the wobble position.

The biosynthesis of s^2^ and xm^5^U modifications requires distinct enzyme sets. In *E. coli*, IscS, TusA-E, and MnmA install the s^2^U modification, whereas in *B. subtilis*, YrvO and MnmA fulfill this role (Figure 1F, **Table 1**) (6). The MnmE-MnmG complex installs xm^5^U modifications (cmnm^5^ with glycine or nm^5^ with NH_4_^+^ as the substrate) in both *E. coli* and *B. subtilis* (16), while MnmC in *E. coli* and MnmL/MnmM in *B. subtilis* provide further modifications (14, 29, 30) (Figure 1). Through comparative genomics and localization studies, we identified the apicoplast orthologs of SufS, MnmA, MnmE, and MnmG in *P. falciparum* (**Table 2**, Figure 3). Our prior studies demonstrated that SufS and MnmA participate in s^2^U modification in the apicoplast (Figure 5) (40) in a manner similar to that observed in *B. subtilis* (6).

**Figure 5.**
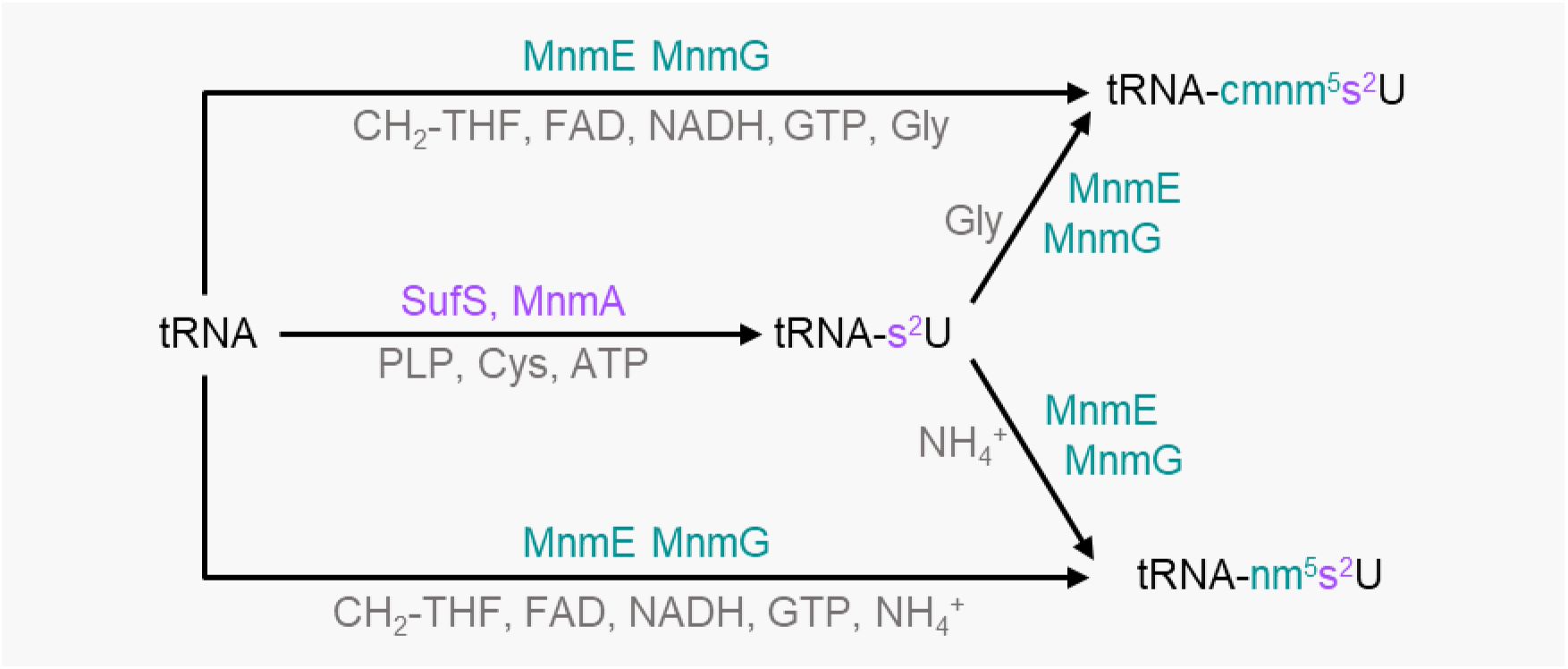
Proposed biosynthesis pathway for xm^5^s^2^U tRNA modifications in *Plasmodium falciparum*. Enzymes responsible for xm^5^s^2^U modification are shown, with respective cofactors and substrates in gray. The MnmE-MnmG complex functions either independently to install xm^5^U modifications (teal) in tRNAs or modifies pre-existing s^2^U bases to generate xm^5^s^2^U. The s^2^U modifications are installed by SufS and MnmA (lavender). No orthologs of MnmC, MnmL, or MnmM have been detected in *P. falciparum*.

MnmA is a highly conserved enzyme and remains in the most reduced bacterial genomes (59–62). Despite extensive searches, however, we could not identify MnmC, MnmL, or MnmM orthologs in malaria parasites. Some organisms, like *Aquifex aeolicus*, encode only MnmC2 and lack MnmC1 because their MnmE-MnmG complex directly produces nm^5^s^2^U using NH_4_^+^ (63). In others, such as *Streptococcus pneumoniae*, only MnmM is present (14). Previous phylogenetic analyses reveal alternative enzyme pairings, like MnmC1/MnmM or MnmL/MnmC2, in other organisms (14, 30). Meanwhile, bacteria with reduced genomes have lost MnmC, MnmL, and MnmM orthologs entirely (59, 64, 65). The conservation of MnmA, MnmE, and MnmG is thought to result in the retention of SufS or other cysteine desulfurases, with some organisms using a dedicated cysteine desulfurase specifically for s^2^U modification in tRNAs (6, 62, 66).

Based on our findings, we propose that the apicoplast of *P. falciparum* contains a minimal xm^5^s^2^U biosynthesis pathway composed of SufS, MnmA/E/G, and that all of these proteins are essential for apicoplast maintenance and parasite survival. As shown in Figure 5, the expected activities of these enzymes require cofactors, including PLP, CH_2_-THF, FAD, SAM, and substrates such as glycine or NH_4_^+^, to be present in the apicoplast. The mechanisms for the synthesis, recycling and/or import of these molecules are largely unknown.

In gram negative bacteria, such as *E. coli* and *Salmonella enterica*, deletion of MnmA results in a slow-growth phenotype in rich media, attributed to the complete absence of s^2^U modifications and an enrichment of xm^5^U modifications in tRNAs (7, 12, 15, 57, 67). In gram positive *B. subtilis*, MnmA deletion is nonviable, suggesting its essential role for survival (6). In *P. falciparum* PfMev lines, deletion of the apicoplast-localized MnmA leads to apicoplast disruption and ultimately parasite death. This phenotype can be reversed by complementation with bacterial MnmA, presumably because a functional MnmA enzyme is required for proper decoding of apicoplast-encoded proteins (40). Deletion of *Toxoplasma gondii* MnmA also results in apicoplast defects and parasite death (68).

MnmE and MnmG form a complex responsible for xm^5^U biosynthesis at the C5 position of tRNAs, and deletions or mutations in these genes significantly impact phenotypic traits. Initial studies in *E. coli* showed that Δ*mnmE* strains exhibit slow growth or lethality, depending on genetic background (17, 18). Similar slow-growth phenotypes are observed in other species, including *B. subtilis* and *Streptococcus suis* (57, 69, 70). The slow growth phenotype is attributed to xm^5^U depletion and s^2^U accumulation in tRNAs following MnmE deletions or mutations (17, 21, 57, 69, 71). The mitochondrial ortholog of MnmE, MSS1 in *Saccharomyces cerevisiae*, also exhibits s^2^U accumulation and reduced oxygen consumption upon deletion, likely from impaired mitochondrial protein translation (72). Likewise, MnmG deletions result in growth impairment or lethality, attributed to xm^5^U depletion (16, 21, 25, 73). Deletion of the *S. cerevisiae* ortholog of MnmG (MTO1) mimics the MSS1 deletion phenotype, consistent with both proteins being required for tRNA modification (72). Both MnmE and MnmG mutations also affect stress responses and pathogenicity across bacterial species (70, 74–77). Interestingly, double deletion of MnmE and MnmG in *S. suis* results in a slow-growth phenotype (78), whereas double deletions of MnmA/MnmE in *S. enterica* or MTU1 (MnmA ortholog)/MSS1 or MTU1/MTO1 in *S. cerevisiae* are lethal (57, 72). These findings show that some organisms can survive without xm^5^U, but not if s^2^U is also missing. In *P. falciparum*, both Δ*mnmE* and Δ*mnmG* lines show apicoplast disruption—loss of the apicoplast genome and formation of vesicles containing nucleus-encoded apicoplast proteins (Figure 4)—paralleling the Δ*mnmA and* Δ*sufS* phenotypes (40) and suggesting that both s^2^U and xm^5^U modifications are essential for apicoplast protein translation.

Recent work identified 28 distinct tRNA modifications from a total tRNA pool of *P. falciparum* (79). Interestingly, the study did not detect any apicoplast-specific tRNA modifications, which the authors attributed to the relatively low contribution of apicoplast-encoded RNAs compared to nuclear-encoded RNAs (0.5-2% rRNA compared to nuclear-encoded rRNAs) (79). However, it is likely that essential tRNA modifications of prokaryotic origin like xm^5^s^2^U do occur in the apicoplast, given the prokaryotic origin of the organelle (33, 34, 36). Methods to selectively enrich apicoplast tRNAs might allow these modifications to be observed in wild-type parasites. Unfortunately, deletion of any enzyme required for xm^5^s^2^U modification results in loss of the apicoplast genome and all apicoplast tRNAs, limiting the value of analyzing mutant strains.

In this study and prior work (40), we identified the enzymes required for generating xm^5^s^2^U tRNA modifications in the *P. falciparum* apicoplast. Together, these findings demonstrate that *P. falciparum* possesses an xm^5^s^2^U biosynthetic pathway (Figure 5) similar to those found in bacteria with highly reduced genomes (59, 64, 65). Based on their phylogenetic conservation, SufS and MnmA/E/G have been included in all proposed minimal genome concepts to date (60, 61, 80–83) and it appears that the *P. falciparum* apicoplast follows this trend. Despite the highly reduced genome and minimal translation system found in the apicoplast organelle, the xm^5^s^2^U biosynthetic pathway has been retained as an indispensable component.

## Materials and Methods

### Bioinformatics

Multiple sequence alignments of MnmE and MnmG orthologs were performed using the ClustalW program in MEGA11 software (84). The alignment included full-length protein sequences from various species: *P. falciparum* (*Pf*MnmE, PlasmoDB ID: PF3D7_0817100; *Pf* MnmG, PlasmoDB ID: PF3D7_1244000), *E. coli* (*Ec* MnmE, UniProt entry: P25522; *Ec* MnmG, UniProt entry: P0A6U3), *B. subtilis* (*Bs* MnmE, UniProt entry: P25811; *Bs* MnmG, UniProt entry: P25812), *Pseudomonas syringae* (*Ps* MnmE, UniProt entry: Q87TS2), *Thermotoga maritima* (*Tm* MnmE, UniProt entry: Q9WYA4; *Tm* MnmG, UniProt entry: Q9WYA1), *Chlorobaculum tepidum* (*Ct* MnmG, UniProt entry: Q8KA85), *Saccharomyces cerevisiae* (*Sc* MSS1, UniProt entry: P32559; *Sc* MTO1, UniProt entry: P53070), *Arabidopsis thaliana* (*At* TrmE, UniProt entry: Q66GQ1; *At* MnmG, UniProt entry: A0A7G2E660), *Homo sapiens* (*Hs* GTPB3, UniProt entry: Q969Y2; *Hs* MTO1, Uniprot entry: Q9Y2Z2),. Sequence alignment output was subsequently visualized using Boxshade (https://junli.netlify.app/apps/boxshade/) to highlight conserved residues across species.

The crystal structure of *T. maritima* MnmE (PDB id: 1XZQ) and *E. coli* MnmG (PDB id: 3GO5) were obtained from RCSB PDB (https://www.rcsb.org/). Homology models of *P. falciparum* MnmE (166-830 amino acid) and MnmG (84-972 amino acid) were predicted with ColabFold v1.5.5 (https://colab.research.google.com/github/sokrypton/ColabFold/blob/main/batch/AlphaF old2_batch.ipynb) (50–52). The highest-ranked predicted models are presented in Figure 2. Visualization of the crystal structures and homology models was performed with UCSF ChimeraX (version 1.5; https://www.rbvi.ucsf.edu/chimerax) molecular visualization program (85). Structural alignments were generated with the ‘*Matchmaker*’ tool of UCSF ChimeraX.

### *Plasmodium falciparum* parental parasite line

To generate knockout and knock-in lines, we used two parental lines: PfMev and PfMev^attB^, respectively (42, 53). Both lines contain an engineered cytosolic mevalonate pathway enabling isoprenoid precursor biosynthesis independent of the apicoplast when mevalonate is present. PfMev^attB^ also includes an attB site in the P230p locus (53). In both lines, the apicoplast is also labeled with a codon-optimized super-folder green fluorescent protein (api-SFG) that enables organelle visualization (42, 53, 55).

### Asexual blood stage culture

Asexual-stage *P. falciparum* parasites were maintained in complete RPMI 1640 medium (CMA) with L-glutamine (USBiological, MA, USA) supplemented with 12.5 μg/mL hypoxanthine, 20 mM HEPES, 0.2% sodium bicarbonate, 5 g/L Albumax II (Life Technologies, CA, USA), and 25 μg/mL gentamicin. Parasites were cultured in human O^+^ red blood cells (RBCs) at 2% hematocrit in 25 cm^2^ tissue culture flasks and were maintained at 37°C under controlled atmospheric conditions (94% N_2_, 3% O_2_, and 3% CO_2_). Parasitemia was monitored every other day via Giemsa-stained thin blood smears and were passaged by diluting infected RBCs with fresh complete medium and uninfected RBCs to ensure continuous propagation.

### Construction of transfection plasmids

The nucleotide sequences encoding the first 165 amino acid residues of MnmE (*mnmE_lp_*) and the first 83 amino acid residues of the MnmG leader peptide (*mnmG_lp_*) were amplified from PfMev^attB^ genomic DNA using specific primers (see **Supplementary Table 3** for primer sequences). Following amplification, the products were digested with *Avr*II and *Bsi*WI restriction enzymes and cloned via ligase-dependent cloning into the corresponding restriction sites of the p15-api-EcDPCK-mCherry vector (53) to produce p15-*mnmE/G_lp_-mCherry* plasmids (**Supplementary Figure 3**).

CRISPR-Cas9 mediated targeted deletion of the *mnmE* (PF3D7_ 0817100) and *mnmG* (PF3D7_1244000) genes was achieved using two plasmids, pRSng (42) and pCasG-LacZ (86) (**Supplementary Figure 4**). For constructing repair plasmids (pRSng-*mnmE/G*), homology arms (HAs; *mnmE* HA1: 488 bp; *mnmE* HA2: 448 bp; *mnmG* HA1: 419 bp; *mnmG* HA2: 424 bp) were amplified from PfMev genomic DNA using HA1 and HA2-specific primers (see **Supplementary Table 3**). These HA1 and HA2 fragments were inserted into the *Not*I and *Ngo*MIV sites, respectively, of the repair plasmids through In-Fusion ligation-independent cloning (LIC, Clontech Laboratories, CA, USA). The gRNA sequences (**Supplementary Table 3**) were inserted into the *Bsa*I sites of the pCasG-LacZ plasmids using LIC as annealed oligonucleotides to generate pCasG-*mnmE/G* gRNA plasmids. All restriction enzymes were obtained from New England Biolabs Inc, MA, USA. Each construct was verified by Sanger sequencing to ensure sequence fidelity.

### Parasite transfections

To generate the *mnmE_lp_^+^* and *mnmG_lp_^+^* parasite lines, PfMev^attB^ parasites were co-transfected with either the p15-*mnmE_lp_-mCherry* or p15-*mnmG_lp_-mCherry* plasmids along with the pINT plasmid (87) encoding mycobacteriophage integrase (75 µg each) using a standard transfection protocol (54). The integrase facilitates targeted integration by recombining the attP site in the expression constructs with the attB site in the parasite genome (**Supplementary Figure 3**). For transfection, 400 µL of RBCs were electroporated with the plasmid mixture at low voltage, then combined with 1.5 mL of PfMev^attB^ cultures (mostly mid-trophozoite to early schizont stages) and incubated in CMA for 48 h. Parasites with successful integration at the P230p attB locus were selected by supplementing the culture medium with 2.5 nM WR99210 (Jacobus Pharmaceuticals, NJ, USA) for seven days. After initial selection, the cultures were maintained in CMA until parasite emergence. Once the parasites appeared, the cultures were maintained in CMA with 2.5 nM WR99210.

For the Δ*mnmE* and Δ*mnmG* deletion lines, RBCs were electroporated with either pRSng-*mnmE* or pRSng-*mnmG* and the respective pCasG plasmids. The transfected RBCs were infected with PfMev parasites and incubated in CMA supplemented with 50 µM mevalonate (Racemic mevalonolactone; Catalog #M4667, Sigma-Aldrich, MO, USA) for 48 h. Selection of parasites with homologous recombination at the native *mnmE* and *mnmG* loci was performed by adding 2.5 nM WR99210 and 50 µM mevalonate to the medium for seven days. Cultures were then maintained in CMA with 50 µM mevalonate until parasites appeared, after which they were cultured in CMA with both 2.5 nM WR99210 and 50 µM mevalonate to sustain the deletion lines.

### Genotype confirmation

Parasite lysates were generated from both parental and transgenic lines by incubating the samples at 90°C for 5 minutes. These lysates served as templates for all subsequent genotype confirmation PCR assays. To validate the genotypes of *mnmE_lp_*^+^ and *mnmG_lp_*^+^, we used specific primer pairs (sequences listed in **Supplementary Table 3**) to amplify the recombinant attL (labeled L) and attR (R) junctions flanking the integrated plasmids. As a control, we also amplified the unmodified attB site (B) from the parental parasite genome. The anticipated sizes of the PCR products are detailed in **Supplementary** Figure 3.

For confirmation of the Δ*mnmE* and Δ*mnmG* genotypes, we employed primer pairs (**Supplementary Table 3**) to target and amplify the 5’ and 3’ regions of the disrupted *mnmE* or *mnmG* loci (referred as Δ5’ and Δ3’, respectively), as well as the corresponding regions from the native loci (labeled 5’ and 3’). The expected sizes of these amplicons are provided in **Supplementary** Figure 4.

### Confirmation of apicoplast genome loss

The presence of the apicoplast genome was assessed by PCR amplification of the apicoplast genome-encoded *sufB* gene (PF3D7_API04700). As controls, we simultaneously amplified the nuclear genome-encoded lactate dehydrogenase gene (*ldh*, PF3D7_1324900) and the mitochondrial genome-encoded cytochrome c oxidase subunit 1 gene (*cox1*, PF3D7_MIT02100). PCR reactions yielded expected amplicon sizes of 520 bp (*ldh*), 581 bp (*sufB*), and 761 bp (*cox1*). All primer sequences are listed in **Supplementary Table 3**. The parental PfMev line served as a positive control for apicoplast genome detection.

### Live epifluorescence microscopy

For live cell imaging of the Δ*mnmE* and Δ*mnmE* transgenic lines, 100 µL of asynchronous parasites were incubated with 1 µg/mL DAPI (62248, Thermo Fisher Scientific, MA, USA) and 30 nM MitoTracker CMX-Ros (M7512, Thermo Fisher Scientific, MA, USA) for 30 min at 37°C. The stained cells were then washed three times with CMA and incubated for 5 min at 37°C after each wash. After the final wash, the cells were resuspended in 20 µL of CMA, placed on a slide and sealed under a cover slip with wax. For live cell imaging of the *mnmE_lp_^+^*and *mnmG_lp_^+^*parasite lines, cells were stained with 1 µg/mL DAPI only.

All images were taken with a Zeiss AxioImager M2 microscope (Carl Zeiss Microscopy, LLC, NY, USA) equipped with a Hamamatsu ORCA-R2 camera (Hamamatsu Photonics, Hamamatsu, Japan) using a 100×/1.4 NA lens. A series of images were taken spanning 5 µm along the z-plane with 0.2 µm spacing. An iterative restoration algorithm using the Volocity software (PerkinElmer, MA, USA) was used to deconvolve the images to report a single image in the z-plane.

### Parasite growth assay

Parasite growth was monitored using an Attune Nxt Flow Cytometer (Thermo Fisher Scientific, MA, USA) following established protocols (42, 88). To assess growth dependence on mevalonate, as shown in Figures 4E and **4I**, Δ*mnmE* and Δ*mnmG* parasites were cultured in the presence or absence of 50 µM mevalonate. Cultures were seeded at an initial parasitemia of 0.5% and a hematocrit of 2%, with a total volume of 250 µL per well, in quadruplicate for each condition. Parasite growth was assessed every 24 hours over a four-day period using SYBR Green I (Catalogue # S7563, Thermo Fisher Scientific, MA, USA) staining. Data from two independent biological replicates (each in quadruplicate) were plotted with Prism V8.4 (GraphPad Software, CA, USA).

## Acknowledgements

This study was supported by the National Institutes of Health (R01 AI125534 to S.T.P.), the Johns Hopkins Malaria Research Institute, and Bloomberg Philanthropies. R.E. received support from the Johns Hopkins Malaria Research Institute postdoctoral fellowship and the Samuel Jordan Graham postdoctoral fellowship. We also acknowledge VEuPathDB, including its curators and contributors, for providing essential data resources that greatly facilitated this research. The funders played no role in study design, data collection and analysis, decision to publish, or manuscript preparation.

## Author contributions

R.E., R.P.S., and S.T.P. conceptualized this work. R.E. wrote the original draft. S.T.P. coordinated this work. R.E., L.R.D., R.P.S., and H.B.L. carried out the experiments. All authors contributed to revision of the manuscript.

