## Supplementary Figures and Tables for "tRNA modifying enzymes MnmE and MnmG are essential for *Plasmodium falciparum* apicoplast maintenance"

```
Pf MnmE     1 MIKRPFVHFVLLKCITVTLFLFIIFVDTFKGKRRNKPKRFPFFIPLKIMKCRKEMHLIKSKIGNRKMFEQKENILSLNME
Ec MnmE     1 --------------------------------------------------------------------------------
Bs MnmE     1 --------------------------------------------------------------------------------
Ps MnmE     1 --------------------------------------------------------------------------------
Tm MnmE     1 --------------------------------------------------------------------------------
Sc MSS1     1 --------------------------------------------------------------------------------
At TrmE     1 ---------------------------------------------------------------------MVSLLCFRSVG
Hs GTB3     1 --------------------------------------------------------------------------------

Pf MnmE    81 EESFIDKNEQNKECEKKNMDINTIHNFLVNNNFASNEDEKKKNICDNISTYYSEANNNMINYVTNYDDVRKEYLLKIMSD
Ec MnmE     1 ----------------------------------------------------------------------------MSD-
Bs MnmE     1 -------------------------------------------------------------------------------M
Ps MnmE     1 ----------------------------------------------------------------------------MNVP
Tm MnmE     1 -------------------------------------------------------------------------------M
Sc MSS1     1 ---------------------------------------------MNSASFLQSRLISRSFLVRRSLKRYSGLAKPYTFQ
At TrmE    12 VYLLRQATPIITRGGRISRPISKNLCFVLFSSSPKTSLLKPRAQSSENNSLVFKGDERVVGLVGKVVDDAFDKVDRFQSS
Hs GTB3     1 ---------------------------------------------MWRGLWTLAAQAAR---GPRRLCTRRSSGAPAPGS

Pf MnmE   161 KETIYALSS-GNVQSAISVIRISGPLSKMILEILLHKGKDKKIKINNNNNNNNNNKNNICYNNYGKRQHFKKDNINRICN
Ec MnmE     4 NDTIVAQAT-PPGRGGVGILRISGFKAREVAETVLG--KLPKPR---------------------------YADYLPFKD
Bs MnmE     2 DTIAAISTPMG--EGAIAIVRLSGPEAIQIADKIYKGPKGKTLSSVES----------------------HTIHYGHIVD
Ps MnmE     5 RETIAAIAT-AQGRGGVGIVRVSGPLAGKTAQAITG--RMPKPR---------------------------FAHYGPFAD
Tm MnmE     2 DTIVAVATPPG--KGAIAILRLSGPDSWKIVQKHLRTRSKIVPR---------------------------KAIHGWIHE
Sc MSS1    36 QPTIYALSTPANQTSAIAIIRISGTHAKYIYNRLVDSSTVPPIR---------------------------KAILRNIYS
At TrmE    92 STIVAIVTPIGGPPGAVGIVRLSGPKAVEVARRVFRSAKKTKKKESDSDT---------------WRPKSHFVEYGAVVD
Hs GTB3    33 GATIFALSS-GQGRCGIAVIRTSGPASGHALRILTAPRDLPLAR---------------------------HASLRLLSD

Pf MnmE   240 EKRTTNDVYNLSYETSGNEKLGDYNLHDIIHMRKNLEERKLYMGKLYDKNNDIIDIVMYSYFKEPKSYTGEELVEIYCHG
Ec MnmE    54 A------------------------------------------------DGSVLDQGIALWFPGPNSFTGEDVLELQGHG
Bs MnmE    58 R-----------------------------------------------PSDRVVEEVMVSVLKAPRTFTREDVIEINCHG
Ps MnmE    55 E------------------------------------------------SGQVLDEGIALYFPGPNSFTGEDVLELQGHG
Tm MnmE    53 -------------------------------------------------NGEDVDEVVVVFYKSPKSYTGEDMVEVMCHG
Sc MSS1    89 PSSCSVKPH------------------------------------DQKESKILLDTSLLLYFQAPYSFTGEDVLELHVHG
At TrmE   157 ------------------------------------------------SNGNVVDEVLAVPMLAPRSYTREDVVELQCHG
Hs GTB3    85 P-----------------------------------------------RSGEPLDRALVLWFPGPQSFTGEDCVEFHVHG

Pf MnmE   320 NMLIVKEIMSAINNMNELFYKIIIEERQNNIYNNDMLCSFIEDNNNNNNNNTKYDTNEGNNNPTNDHIIYEKIHERCHHN
Ec MnmE    86 GPVILDLLLKRILTIPG---------------------------------------------------------------
Bs MnmE    91 GIVTVNQVLQLALREGA---------------------------------------------------------------
Ps MnmE    87 GPIVLDMLLQRCLQL-G---------------------------------------------------------------
Tm MnmE    84 GPLVVKKLLDLFLKSGA---------------------------------------------------------------
Sc MSS1   133 GKAVVNSILKAIGSLHDRSS------------------------------------------------------------
At TrmE   189 SEVCLRRVLRTCVEAGA---------------------------------------------------------------
Hs GTB3   118 GPAVVSGVLQALGSVPG---------------------------------------------------------------

Pf MnmE   400 FIKIRESYKGEFTRRAFENNKMSLLQIEGLKELLFCKQKVQKQIALNYMNGYAKDIYLKLKELLKELLVYTELKIDFEEE
Ec MnmE   103 ---LRIARPGEFSERAFLNDKLDLAQAEAIADLIDASSEQAARSALNSLQGAFSARVNHLVEALTHLRIYVEAAIDFPDE
Bs MnmE   108 ----RLAEPGEFTKRAFLNGRIDLSQAEAVMDLIRAKTDRAMNVAMNQMEGRLSALVRRLRSEILETLAHVEVNIDYPEY
Ps MnmE   103 ---SRLARPGEFSERAFLNDKLDLAQAEAIADLIEASSAQAARNALRSLQGVFSQRVDNLTEKLISLRIYVEAAIDFPEE
Tm MnmE   101 ----RMAEPGEFTKRAFLNGKMDLTSAEAVRDLIEAKSETSLKLSLRNLKGGLRDFVDSLRRELIEVLAEIRVELDYP--
Sc MSS1   153 GKDIRFALPGDFSRRAFQNGKFDLTQLEGIKDLIDSETESQRRSALSSFNGDNKILFENWRETIIENMAQLTAIIDFADD
At TrmE   206 ----RLAEPGEFTLRAFLNGRLDLSQAENVEKLISAKSSAAADAALEGIQGGFSSLVKSLRAQCIELLTEIEARLDF--E
Hs GTB3   135 ---LRPAEAGEFTRRAFANGKLNLTEVEGLADLIHAETEAQRRQALRQLDGELGHLCRGWAETLTKALAHVEAYIDFGED

Pf MnmE   480 HITSPKEKEEIQNMVLRKIREAINHINYILKKNNVEDISNS-FDILLFGNVNSGKSTLMNNICNNDISIVTNIKGSTLDI
Ec MnmE   180 EIDFLSD-----GKIEAQLNDVIADLDAVRAEARQGSLLREGMKVVIAGRPNAGKSSLLNALAGREAAIVTDIAGTTRDV
Bs MnmE   184 DDVEEMT----HQILVEKATAVKKEIETLLRTSEQGKILREGLSTVIIGRPNVGKSSLLNSLVHEAKAIVTDIPGTTRDV
Ps MnmE   180 EIDFLAD-----GHVLGMLDDVRAELSTVLREAGQGALLRDGMTVVIAGRPNAGKSSLLNALAGREAAIVTEIAGTTRDV
Tm MnmE   175 DEIETN-----TGEVVTRLERIKEKLTEELKKADAGILLNRGLRMVIVGKPNVGKSTLLNRLLNEDRAIVTDIPGTTRDV
Sc MSS1   233 NSQEIQNTDEIFHNVEKNIICLRDQIVTFMQKVEKSTILQNGIKLVLLGAPNVGKSSLVNSLTNDDISIVSDIPGTTRDS
At TrmE   280 DEMPPLD----IESVINKITSMSQDVESALDTANYDKLLQSGLQIAIVGRPNVGKSSLLNAWSKSERAIVTEVAGTTRDV
Hs GTB3   212 DNLEEG----VLEQADIEVRALQVALGAHLRDARRGQRLRSGVHVVVTGPPNAGKSSLVNLLSRKPVSIVSPEPGTTRDV

Pf MnmE   559 IQKNIQIFNNSYNLCDSAGVIKSEKIKSLNEKNYILYEKKKKKKKNIHKTLESMGIKKTLSFLKNNCISAFVLLNIKNYK
Ec MnmE   255 LREHIHIDGMPLHIIDTAGLR------------------------EASDEVERIGIERAWQEIEQADRVLFMVDGTTTDA
Bs MnmE   260 IEEYVNVRGVPLRLVDTAGIR------------------------ETEDIVERIGVERSRQVLKEADLILLVLNYSEELS
Ps MnmE   255 LREHIHIDGMPLHVVDTAGLR------------------------DTQDQVEMIGVQRALKAIGEADRILLVVDATAPEA
Tm MnmE   250 ISEEIVIRGILFRIVDTAGVRS-----------------------ETNDLVERLGIERTLQEIEKADIVLFVLDASSPLD
Sc MSS1   313 IDAMINVNGYKVIICDTAGIRE-----------------------KSSDKIEMLGIDRAKKKSVQSDLCLFIVDPTDLSK
At TrmE   356 VEANVTVRGVPITLLDTAGIR------------------------ETNDIVEKIGVERSETAAKVADVIIMAVSAVEGWT
Hs GTB3   288 LETPVDLAGFPVLLSDTAGLRE-----------------------GVG-PVEQEGVRRARERLEQADLILAMLDASDLAS
```

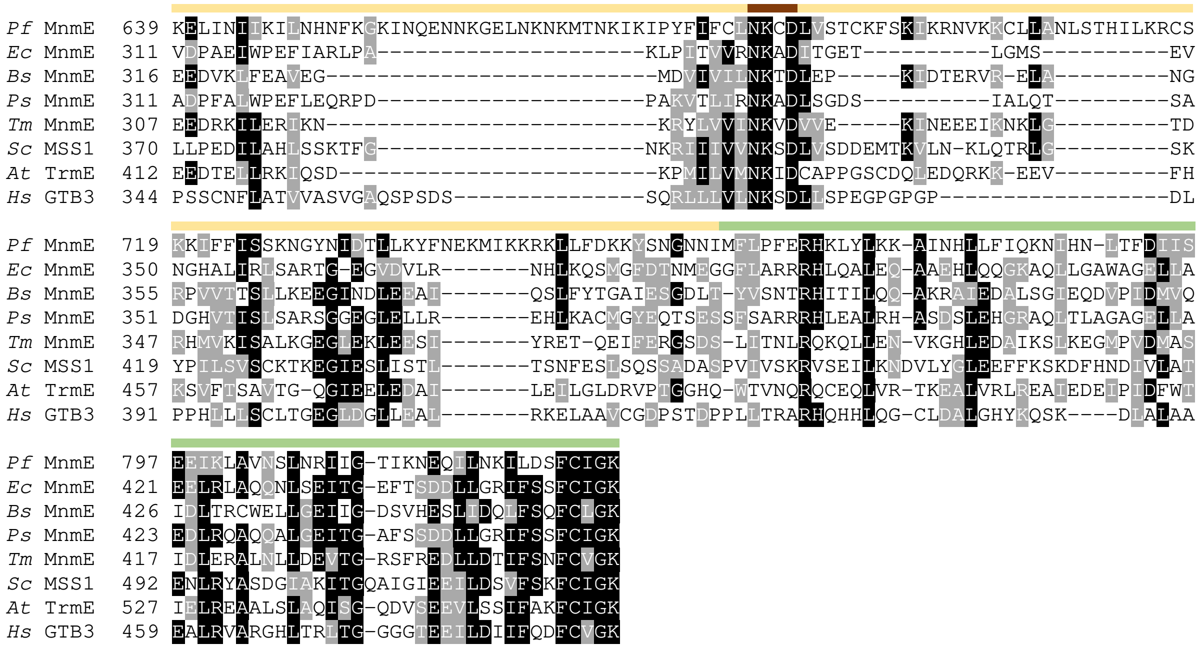


**Supplementary Figure 1.**

Multiple sequence alignment (MSA) of MnmE orthologs from *Plasmodium falciparum* (*Pf* MnmE), *Escherichia coli* (*Ec* MnmE), *Bacillus subtilis* (*Bs* MnmE), *Pseudomonas syringae* (*Ps* MnmE), *Thermotoga maritima* (*Tm* MnmE), *Saccharomyces cerevisiae* (*Sc* MSS1), *Arabidopsis thaliana* (*At* TrmE), and *Homo sapiens* (*Hs* GTB3). MSA output was visualized using Boxshade to highlight conserved residues. Amino acid residues and domains important for MnmE activity are depicted as in **Figure 2A**.

```
PfMnmG    1 MFIIIIFFEMFIRFNKFLKLKVSFYSLLLLLLFIIYIFFFCSPCYAYNFSKVKQTFLWPYSPPKIKRGNHLKSSQISEKC
EcMnmG    1 --------------------------------------------------------------------------------
BsMnmG    1 ------------------------------------------------------------------------------MG
TmMnmG    1 -----------------------------------------------------------------------------MIG
CtMnmG    1 --------------------------------------------------------------------------------
AtMnmG    1 ----------------MRAVAAATATVSLRHFRSFPTIVPCLLFSSSSSLPFHSPRLCVFLRPRQLFLNRPLAASFSSSS
ScMTO1    1 ---------------------------------------------------MLRVTTLAS-SCTSFPLQVLRRRLTISSL
HsMTO1    1 ---------------------------------------------------MFYFRGCGRWVAVSFTKQQFP-LARLSSD

PfMnmG   81 PCNNVKKNYDVIVIGGGHSGCEASYISAKSNAMTLLVTQNKETIGEMSCNPSIGGIGKGILVKEIDALGGLMGKVIDKSG
EcMnmG    1 --MFYPDPFDVIIIGGGHAGTEAAMAAARMGQQTLLLTHNIDTLGQMSCNPAIGGIGKGHLVKEVDALGGLMAKAIDQAG
BsMnmG    3 YEAG---QYDVIVIGAGHAGVEAALASARQGAKTLVLTINLDMVAFMPCNPSVGGPAKGIVVREIDALGGEMGRNIDKTH
TmMnmG    4 LRPEDDRVYDVIVVGAGHAGIEAALAAARMGFRVLVLTVNPDTVGWAPCNPAIGGPAKGVVVREIDALGGEMAKTTDETM
CtMnmG    1 -------MYDVIVVGAGHAGCEAALAVARGGLHCLLITSDLSAVARMSCNPAIGGVAKGQITREIDALGGEMGKAIDATG
AtMnmG   65 SGATSDSTYDVIVVGAGHAGCEAALASARLGASTLLLTLNLDRIAWQPCNPAVGGPAKSQLVHEVDALGGDIGKVADRCY
ScMTO1   29 TSFQPTTKTQVVVIGAGHAGCEAAAASSRTGAHTTLITPSLTDIGKCSCNPSIGGVGKGILVKEIDALDGLMGKVTDLAG
HsMTO1   29 SAAPRTPHFDVIVIGGGHAGTEAATAAARCGSRTLLLTHRVDTIGQMSCNPSFGGIGKGHLMREVDALDGLCSRICDQSG

PfMnmG  161 IHFKILNMKKGLAVRGHRAQADRDLYNYYMKEYLYN---TPNLDILENSVQSLLIDTYEDKNNNNDNINDYNNNVKKKKI
EcMnmG   79 IQFRILNASKGPAVRATRAQADRVLYRQAVRTALEN---QPNLMIFQQAVEDLIVE--------------------NDRV
BsMnmG   80 IQMRMLNTGKGPAVRALRAQADKFQYQHEMKNTLEK---EPNLTLLQGIVERLIVE--------------------DGEC
TmMnmG   84 INVRMLNVSKGPAVRALRAQIDKISYSRTMKRKLET---NPNIVLRHGIVERILTE--------------------KGRV
CtMnmG   74 IQFRMLNRSKGPAMHSPRAQADKTQYSLYMRRIVEH---EPNIDLLQDTVIGVSAN--------------------SGKF
AtMnmG  145 LQKRILNVSRGPAVRSLRAQTDKREYAMEMKKIVDS---TENLCIREAMVTDIIVGK-------------------NDNV
ScMTO1  109 VQFKMLNRSKGPAVWGPRAQIDRELYKKYMQRELSDKKAHPNLSLLQNKVADLILYDPG---------------CGHKVI
HsMTO1  109 VHYKVLNRRKGPAVWGLRAQIDRKLYKQNMQKEILN---TPLLTVQEGAVEDLILTEPEPEH------------TGKCRV

PfMnmG  238 YGIKNKCSCEIYSHNVILTTGTFLGGICHIGKDKYIGGRIKRLINNFDLKMKKGNIESIKTNDEPIIKQDKLKKINQNGE
EcMnmG  136 VGAVTQMGLKFRAKAVVLTVGTFLDGKIHIGLDNYSGGR-----------------------------------------
BsMnmG  137 RGVITQTGAHYRAKAVVMTTGTYLRGRIILGDLSYSSGP-----------------------------------------
TmMnmG  141 KGVVDNYGIDYLGKAVIVTTGTFLRGKIFIGRSTFPAGR-----------------------------------------
CtMnmG  131 SSVTVRSGRAIQAKAAILACGTFLNGLIHIGMDHFPGGRS----------------------------------------
AtMnmG  203 EGVATFFGMNFYAPSVILTTGTFMSGKIWVGKKSMPAGR-----------------------------------------
ScMTO1  174 KGVVLDDGTQVGADQVIITTGTFLSAEIHIGDKRIAAGR-----------------------------------------
HsMTO1  174 SGVVLVDGSTVYAESVILTTGTFLRGMIVIGLETHPAGR-----------------------------------------

PfMnmG  318 STINNMHNNYNEDNKKYINYEENLNIPFEKLIESSTKNIAQQLR-ENNFEIKRMKTGTPPRLDINSINFHVLEREDTEKD
EcMnmG  175 ------------------------------AGDPPSIPLSRRLR-ELPLRVGRLKTGTPPRIDARTIDFSVLAQQHGDNP
BsMnmG  176 ------------------------------NNQQPSIKLSEHLE-ELGFDLVRFKTGTPPRVKSDTIDYSKTEIQPGDDV
TmMnmG  180 ------------------------------MGEFPATKLTESLI-ELGFEVGRFKTGTPARVLKRSINFSVMERQDTSDE
CtMnmG  171 ------------------------------TAEPPVEGLTESLA-SLGFSFGRLKTGTPPRIDSRSVDYTIVTEQPGDVD
AtMnmG  242 ------------------------------AGESASQGLTENLQ-KLGFETDRLKTGTPARVDRRTIDFSNLEAQHGDEE
ScMTO1  213 ------------------------------IGEQPTYGISNTLQNEVGFQLGRLKTGTPARLAKESIDFSALEVQKGDAL
HsMTO1  213 ------------------------------LGDQPSIGLAQTLE-KLGFVVGRLKTGTPPRIAKESINFSILNKHIPDNP

PfMnmG  397 YPFYFSFLNMNKINN-NKTLPCYKTYTNLKTHELVRNNLAQLPDYDS-FDKFGNGPRYCPSIAKKVMKFSEKKKHIIWIE
EcMnmG  224 -MPVFSFMGNASQHP--QQVPCYITHTNEKTHDVIRSNLDRSPMYA--GVIEGVGPRYCPSIEDKVMRFADRNQHQIFLE
BsMnmG  225 -PRAFS-YETVEYIT--DQLPCWLTYTSPETHEIIDSNLHRSPMYS--GMIKGTGPRYCPSIEDKVVRFNDKPRHQIFLE
TmMnmG  229 -PLAFSFFDEPRVLP--KDYPCWLTRTNPETHSIIKQYLEFSPLYGTVKLIEGIGPRYCPSIEDKVVKFKDKESHQVFVE
CtMnmG  220 -PVPFSFSSTSVANR--NLVSCYLTKTTEKTHDILRTGFDRSPLFT--GKVQGVGPRYCPSIEDKISRFPDKSSHHIFLE
AtMnmG  291 -VSWFSFDPDFHIER--EQMCCYLTRTTKITHQLIRDNLHETPTYG--GWVEAKGPRYCPSIEDKIVRFKDKESHQIFLE
ScMTO1  263 -PVPMSFLNETVSVEPTKQLDCFGTHTTPQMHDFLRNNLHQSIHIQ---DTTIKGPRYCPSIEAKILRFPDRSSHKIWLE
HsMTO1  262 -SIPFSFTNETVWIKPEDQLPCYLTHTNPRVDEIVLKNLHLNSHVK---ETTR-GPRYCPSIESKVLRFPNR-LHQVWLE

PfMnmG  475 PEGFNNKLIYPNGLSSAYPIHIQKDIINSIKGLENAQIVFP-------------------------AYDVEYYYVNPKCL
EcMnmG  299 PEGLTSNEIYPNGISTSLPFDVQMQIVRSMQGMENAKIVRP-------------------------GYAIEYDFFDPRDL
BsMnmG  299 PEGRNTQEVYVQGLSTSLPEDVQQRMLATIPGLENVQMMRA-------------------------GYAIEYDAIVPTQL
TmMnmG  306 PEGRDTEEYYLNGLSTSLPYEAQIKMIRSVKGLENAIVTRP-------------------------AYAIEYDYIDPRQL
CtMnmG  295 PEGTDIVEMYVNGFSTSLPEDIQIAGLRSIPGLEEAKMIRP-------------------------GYAIEYDFFHPWQI
AtMnmG  366 PEGRDVPEIYVQGFSTGLPENLQLPLLRSLPGLENCSMLRP-------------------------AYAVEYDYLPAHQC
ScMTO1  339 PEGFNSDVIYPNGISNSMPEDVQLQMMRLIPGMANVEILQP-------------------------AYGVEYDYVDPRQL
HsMTO1  336 PEGMDSDLIYPQGLSMTLPAELQEKMITCIRGLEKAKVIQPDGVLLLLPRMECNGAISAHHNLPLPGYGVQYDYLDPRQI

PfMnmG  530 NYTLETKNVKGLFLAGQICGTTGYEEAACQGIVAGINAALNSKKDNNKYNNNNNNENNNNHNINNDEVNKFILKRNESYI
EcMnmG  354 KPTLESKFIQGLFFAGQINGTTGYEEAAAQGLLAGLNAARLSAD-----------------------KEGWAPARSQAYL
BsMnmG  354 WPTLETKKITNLYTAGQINGTSGYEEAAGQGIMAGINAGRKALG-----------------------KEEVILSRSDAYI
TmMnmG  361 YPTLESKLVENLYFAGQVNGTSGYEEAAGQGIIAGINAALKLRG-----------------------EPPLILKRSEAYI
CtMnmG  350 RSTMETRPVENLFFAGQINGTSGYEEAAAQGLMAGINAVRKILG-----------------------KELIVLGRDQAYI
AtMnmG  421 SRSLMTKKIEGLFFSGQINGTTGYEEAAAQGIISGINAARHADG-----------------------KKHVVLERESSYI
ScMTO1  394 KPSLETKLVDGLFLAGQINGTTGYEEAAAQGIIAGINAGLLSRQE----------------------REQLVLKRSEAYI
HsMTO1  416 TPSLETHLVQRLFFAGQINGTTGYEEAAAQGVIAGINASLRVSR-----------------------KPPFVVSRTEGYI
```

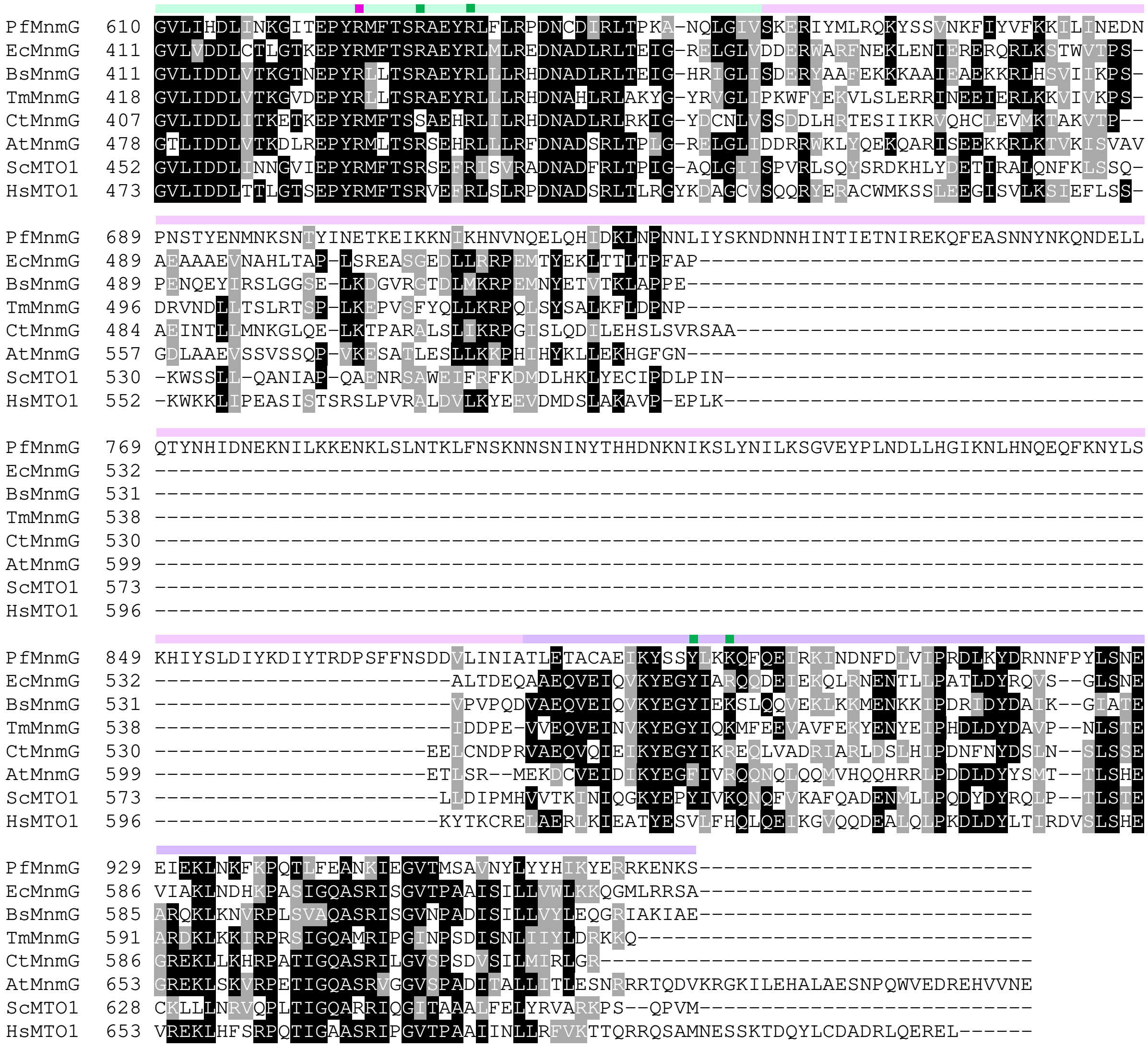


**Supplementary Figure 2.**

Multiple sequence alignment (MSA) of MnmG orthologs from *Plasmodium falciparum* (*Pf* MnmG), *Escherichia coli* (*Ec* MnmG), *Bacillus subtilis* (*Bs* MnmG), *Thermotoga maritima* (*Tm* MnmG), *Chlorobaculum tepidum* (*Ct* MnmG), *Arabidopsis thaliana* (*At* MnmG), *Saccharomyces cerevisiae* (*Sc* MTO1), and *Homo sapiens* (*Hs* MTO1). MSA output was visualized using Boxshade to highlight conserved residues. Amino acid residues and domains important for MnmG activity are depicted as in **Figure 2C**.

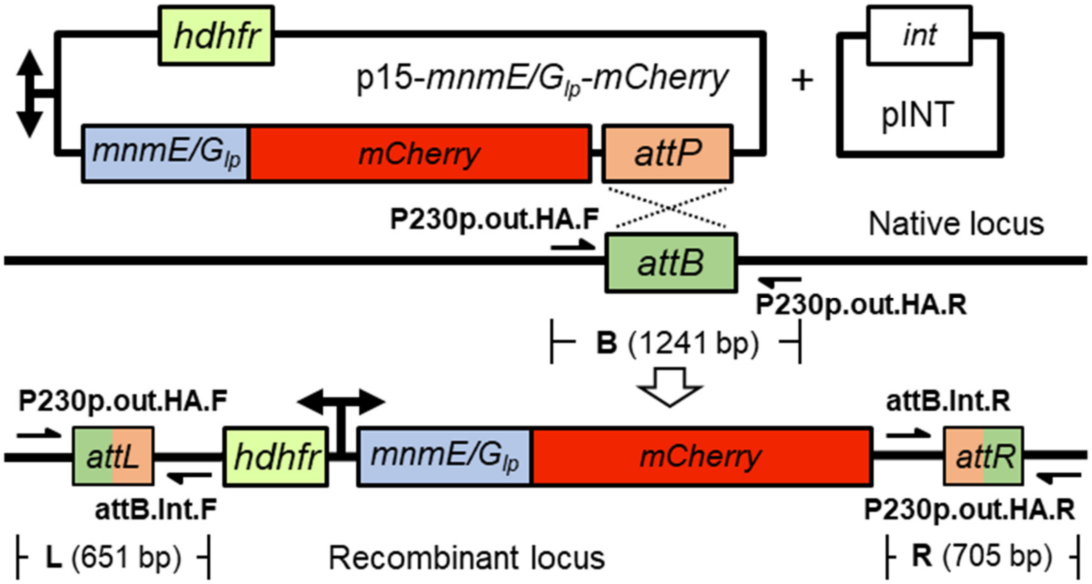


**Supplementary Figure 3.**
Schematic illustration of p15-*mnmE/G*$_{lp}$-mCherry plasmid insertion into the attB locus of PfMev$^{attB}$ parasites to generate the *mnmE/G*$_{lp}^{+}$ line. The *mnmE/G*$_{lp}$*-mCherry* and human dihydrofolate reductase (*hdhfr*) selection cassettes are expressed under a single bidirectional promoter. This plasmid was co-transfected with the pINT plasmid, which expresses integrase (*int*) for catalyzing attB/attP recombination. Primer pairs (black half arrows) for PCR amplification of the attB (B) region of the parental line and the recombinant attL (L) and attR (R) regions are marked along with the expected amplicon sizes. Refer to **Supplementary Table 3** for primer sequences.

**A**

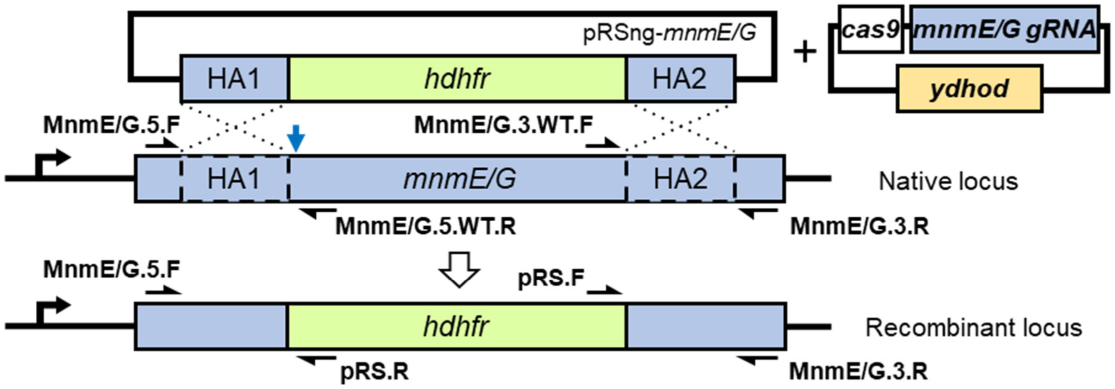


**B**

| PCR reaction | Primer combination | | Parasite line |
|---|---|---|---|
| | Forward | Reverse | |
| Δ5’ | MnmE/G.5.F | pRS.R | Δ*mnmE*/Δ*mnmG* |
| Δ3’ | pRS.F | MnmE/G.3.R | |
| 5’ | MnmE/G.5.F | MnmE/G.5WT.R | |
| 3’ | MnmE/G.3WT.F | MnmE/G.3R | |
| Δ5’ | MnmE/G.5.F | pRS.R | PfMev (parental) |
| Δ3’ | pRS.F | MnmE/G.3.R | |
| 5’ | MnmE/G.5.F | MnmE/G.5WT.R | |
| 3’ | MnmE/G.3WT.F | MnmE/G.3R | |

**C**

| | Anticipated amplicon sizes (bp) | | | |
|---|---|---|---|---|
| Gene of interest | Δ5’ | Δ3’ | 5’ | 3’ |
| *mnmE* | 563 | 553 | 607 | 623 |
| *mnmG* | 628 | 556 | 854 | 610 |

**Supplementary Figure 4.**
(A) Schematic illustrating double-crossover homologous recombination for *mnmE/G* gene knockout. A repair plasmid (pRSng-*mnmE/G*) harboring two homology arms (HAs) flank the desired modification site within the native *mnmE/G* locus. Cas9 endonuclease with guide RNA (Cas9-*mnmE/G gRNA* plasmid) introduces a double-stranded break (blue arrow) in the native locus, facilitating homologous recombination with the repair plasmid leading to the recombinant Δ*mnmE/G* locus. Primer positions and directions (black half arrows) for confirming gene knockout are indicated. Refer to **Supplementary Table 3** for primer sequences. *hdhfr*,

human dihydrofolate reductase; *ydhod*, yeast dihydroorotate dehydrogenase. (B) Table containing the PCR reactions, primer combinations used to confirm integration of plasmid, and the respective parasite line. (C) Table depicting the expected amplicon size in base pairs (bp) of each PCR reaction for each gene of interest.

**Supplementary Table 1.** List of enzymes and corresponding UniProt entries involved in the $xm^5s^2U$ modification biosynthesis pathway in model bacteria.

| Enzyme | UniProt entry | Bacteria species |
|---|---|---|
| IscS | P0A6B7 | *Escherichia coli* |
| YrvO | O34599 | *Bacillus subtilis* |
| TusA | P0A890 | *Escherichia coli* |
| TusB | P45530 | *Escherichia coli* |
| TusC | P45531 | *Escherichia coli* |
| TusD | P45532 | *Escherichia coli* |
| TusE | P0AB18 | *Escherichia coli* |
| MnmA | P25745 | *Escherichia coli* |
| | O35020 | *Bacillus subtilis* |
| MnmC | P77182 | *Escherichia coli* |
| MnmE | P25522 | *Escherichia coli* |
| | P25811 | *Bacillus subtilis* |
| MnmG | P0A6U3 | *Escherichia coli* |
| | P25812 | *Bacillus subtilis* |

**Supplementary Table 2.** Bacterial enzymes and their orthologs in apicomplexan parasites associated with the $xm^5s^2U$ modification biosynthesis pathway. ND, not detected.

| Bacterial enzymes | Apicomplexans | Gene ID | % sequence identity | E-value |
|---|---|---|---|---|
| IscS | *Plasmodium falciparum* | PF3D7_0727200 | 50% | 1e-142 |
| | *Toxoplasma gondii* | TGME49_211090 | 50% | 7e-140 |
| | *Eimeria tenella* | ETH2_1354600 | 49% | 7e-127 |
| | *Babesia microti* | BMR1_03g04230 | 53% | 2e-136 |
| | *Theileria annulata* | TA19715 | 53% | 2e-151 |
| | *Cryptosporidium parvum* | cgd4_3040 | 53% | 8e-145 |
| YrvO | *Plasmodium falciparum* | PF3D7_0716600 | 24% | 3e-13 |
| | *Toxoplasma gondii* | TGME49_216170 | 29% | 1e-13 |
| | *Eimeria tenella* | ETH2_1112500 | 27% | 4e-06 |
| | *Babesia microti* | BMR1_03g01415 | 25% | 1e-21 |
| | *Theileria annulata* | TA16295 | 25% | 7e-15 |
| | *Cryptosporidium parvum* | ND | - | - |
| MnmA | *Plasmodium falciparum* | PF3D7_1019800 | 30% | 2e-13 |
| | *Toxoplasma gondii* | TGME49_309110 | 31% | 2e-36 |
| | *Eimeria tenella* | ETH2_1349800 | 38% | 2e-43 |
| | *Babesia microti* | BMR1_03g03155 | 38% | 1e-59 |
| | *Theileria annulata* | TA12620 | 35% | 7e-54 |
| | *Cryptosporidium parvum* | ND | - | - |
| MnmE | *Plasmodium falciparum* | PF3D7_0817100 | 24% | 8e-23 |
| | *Toxoplasma gondii* | TGME49_204350 | 34% | 3e-18 |
| | *Eimeria tenella* | ETH2_1106900 | 36% | 8e-14 |
| | *Babesia microti* | BMR1_02g02630 | 31% | 2e-59 |
| | *Theileria annulata* | TA17200 | 31% | 4e-39 |
| | *Cryptosporidium parvum* | ND | - | - |
| MnmG | *Plasmodium falciparum* | PF3D7_1244000 | 44% | 6e-85 |
| | *Toxoplasma gondii* | TGME49_242010 | 50% | 2e-59 |
| | *Eimeria tenella* | ETH2_1518800 | 37% | 4e-72 |
| | *Babesia microti* | BmR1_04g09035 | 40% | 1e-118 |
| | *Theileria annulata* | TA09230 | 44% | 3e-134 |
| | *Cryptosporidium parvum* | ND | - | - |

**Supplementary Table 3.** Primers used in this study. Restriction enzyme sites are underlined. Guide RNA sequences are in lower case.

| Primer name | Sequence (5'→3') | Primer description |
|---|---|---|
| Primers to amplify homology arms (HA) and guide RNA (gRNA) annealing for *mnmE* and *mnmG* knockout | | |
| MnmE.HA1F | GCCACGAGCGGCCCTTTGTTCATTTCGTTTTGTTAAAATGC | Forward for HA1 amplification |
| MnmE.HA1R | AAGCGCAGCGGCCTAAGGGCATATATCGTTTCTTTATCACTCATA | Reverse for HA1 amplification |
| MnmE.HA2F | CGACAGACGCCGGAGGGGGAATTGAATAAAAATAAAATGACAAA | Forward for HA2 amplification |
| MnmE.HA2R | GGCCACCAGCCGGCCTATGATACGATTCAAAGAATTGACAGCTAG | Reverse for HA2 amplification |
| MnmE.gRNA.F | TAAGTATATAATATTttgcacattacccgaactaaGTTTTAGAGCTAGAA | guide RNA forward oligo |
| MnmE.gRNA.R | TTCTAGCTCTAAAACttagttcgggtaatgtgcaaAATATTATATACTTA | guide RNA reverse oligo |
| MnmG.HA1F | GCCACGAGCGGCCACATTAGAATATGTAACTCGAGGTATATACAAAC | Forward for HA1 amplification |
| MnmG.HA1R | AAGCGCAGCGGCCGTATGGCCATAAAAATGTCTGCTTTACCTTTG | Reverse for HA1 amplification |
| MnmG.HA2F | CGACAGACGCCGGAAACTAATATAAGGGAAAAACAATTTGAAGCATC | Forward for HA2 amplification |
| MnmG.HA2R | GGCCACCAGCCGGATCTCTGCACAGGCTGTTTCTAATGTTGC | Reverse for HA2 amplification |
| MnmG.gRNA.F | TAAGTATATAATATTttttacttggggggagagtaGTTTTAGAGCTAGAA | guide RNA forward oligo |
| MnmG.gRNA.R | TTCTAGCTCTAAAACtactctccccccaagtaaaaAATATTATATACTTA | guide RNA reverse oligo |
| Primers for gene knockout confirmation | | |
| MnmE.5.F | TGTTTTATCCTTATAATACCTTTATTAAACAACTAGATGA | Forward for 5' and Δ5' PCR |
| MnmE.5.WT.R | CCAATATCATTTTGCTCAACGGACCAG | Reverse for 5' PCR |
| MnmE.3.WT.F | CGGCATTTGTATTATTAAATATTAAAAACTACAAGAAGG | Forward for 3' PCR |
| MnmE.3.R | CCCTTTTAATCACTTTCCTATACAAAAACTATCC | Reverse for 3' and Δ3' PCR |

| | | |
|---|---|---|
| MnmG.5.F | CCTTTCACGTTAGGCGCTATTATTCATATAACTAG | Forward for 5’ and Δ5’ PCR |
| MnmG.5.WT.R | CTGAGATTTGTGAAGATTTTAAATGGTTTCCTC | Reverse for 5’ PCR |
| MnmG.3.WT.F | TGTAAATCAAGAACTACAACATATAGACAAATTAAATCCT | Forward for 3’ PCR |
| MnmG.3.R | CCTGTCATATTTTAAATCTCTAGGAATAACTAAATCAAAG | Reverse for 3’ and Δ3’ PCR |
| pRS.F | CATATTTATTAAATCTAGAATTCGACAGACGCCG | Forward for Δ3’ PCR |
| pRS.R | TACAAAATGCTTAAGCGCAGCGGCC | Reverse for Δ5’ PCR |
| Primers to amplify representative genes from nuclear and organellar genome | | |
| LDH.F | GGAGATGTAGTTTTGTTCGATATTG | Forward for PCR |
| LDH.R | CTTGTAAAGGGATACCACCTACAG | Reverse for PCR |
| SufB.F | CATGTAGCTATAGTAGAAATAATAGTAAAAGATTATGG | Forward for PCR |
| SufB.R | GACTCTGAAATACTTAAACCACGTTGC | Reverse for PCR |
| Cox1.F | CTTCATCTTTAAGAATAATTGCACAAGAAAATGTAAATC | Forward for PCR |
| Cox1.R | GTACATATGATGTACCCATACTAAGCTTCC | Reverse for PCR |
| Primers for generation of p15-*mnmE/G*$_{lp}$-*mcherry* plasmid | | |
| MnmElocF | ACAACCTAGGATGATAAAGAGACCCTTTGTTCATT | Forward for $mnmE_{lp}$ amplification |
| MnmElocR | TGCTCGTACGTCTTATCACACTTATAGCACTTTGC | Reverse for $mnmE_{lp}$ amplification |
| MnmGlocF | ACAACCTAGGATGTTTATAATTATTATTTTTTTTGAAATG | Forward for $mnmG_{lp}$ amplification |
| MnmGlocR | TGCTCGTACGATTTTGAGTTACTAGTAAAGTC | Reverse for $mnmG_{lp}$ amplification |
| Primers for genotyping of $mnmE_{lp}^+$ and $mnmG_{lp}^+$ parasites | | |
| P230p.out.HA.F | GGTTGTGATTTTTCAGGTGATTCC | Forward for L and B PCR |
| P230p.out.HA.R | GAAAATTGTAGGGGCAGCTAAATCCGAC | Reverse for R and B PCR |
| attB.Int.F | GCAGTGTGGAATTCCCTGCA | Reverse for L PCR |
| attB.Int.R | TTAAGTGTAGTTAATTCATCAAATAGCATGC | Forward for R PCR |
| Primers for sequencing | | |

| | | |
|---|---|---|
| pRS.R | TACAAAATGCTTAAGCGCAGCGGCC | For HA1 insertion in pRSng-*mnmE/G* |
| pRS.F | CATATTTATTAAATCTAGAATTCGACAGACGCCG | For HA2 insertion in pRSng-*mnmE/G* |
| pL6.gRNA.F | GGGTAAATTATTATTAAAAAATGTATATGTTATG | For guide RNA insertion in pCasG-LacZ |
| RFP.R | GAGGGCTCCGTGAACGGC | Reverse for *mnmE/G*$_{lp}$ insertion in p15-*mnmE/G*$_{lp}$*-mcherry* |
